# Analysis of the persistence and particle size distributional shift of sperm-derived environmental DNA to monitor Jack Mackerel spawning activity

**DOI:** 10.1101/2022.03.09.483695

**Authors:** Satsuki Tsuji, Hiroaki Murakami, Reiji Masuda

## Abstract

Environmental DNA (eDNA) analysis holds great promises as an efficient and noninvasive method to monitor not only the presence and biomass of organisms but also their spawning activity. In eDNA analysis-based monitoring of spawning activity, the detection of sperm-derived eDNA is a key point; however, its characteristics and dynamics are completely unknown. The present study focuses on the persistence and particle size distribution (PSD) of eDNA derived from the sperm of Japanese jack mackerel. First, we investigated the time-dependent degradation and the PSD of sperm-derived eDNA by artificially adding sperm to fish eDNA-free seawater. Next, we kept fish in tanks and examined the changes in DNA concentration and PSD before and after spawning. The data obtained from the two experiments showed that the degradation of sperm-derived eDNA proceeded rapidly, with PSD shifting to a smaller size, regardless of the DNA region (Cyt *b* or ITS1). Additionally, it was shown that the nuclei and mitochondria released from sperm through degradation had a size distribution that was not simply dependent on each organelle size. The results of this study will contribute to elucidating the characteristics and dynamics of eDNA specifically during the spawning season and to further developing eDNA analysis as a powerful tool for the monitoring of spawning activity.

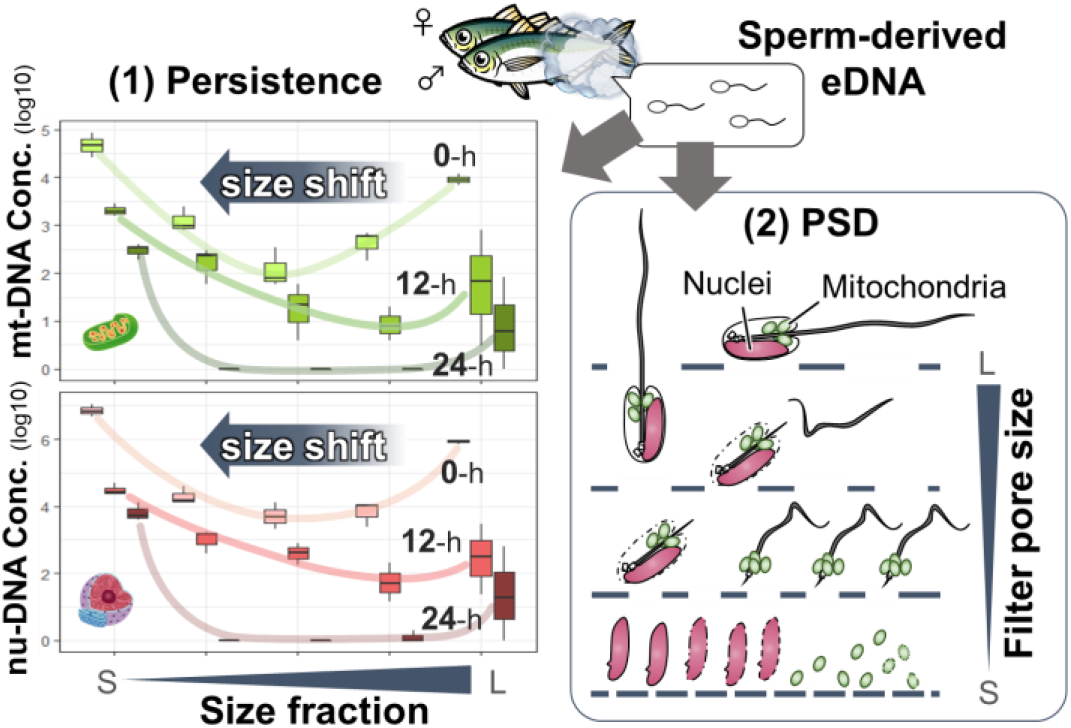

## Introduction

Environmental DNA (eDNA) analysis is an emerging and powerful tool for assessing the diversity, distributions and biomass of organisms ^1–3^. In eDNA analysis, investigators detect the genetic materials shed by organisms into the environment such as skin, faeces, mucus and reproductive materials (e.g., oocytes, ovarian fluid and sperm)^4–7^. The popularity of eDNA analysis has increased rapidly over the past decade, mainly because of its higher cost and time effectiveness, its specificity and sensitivity and its noninvasiveness compared with traditional monitoring methods^8–11^. Moreover, in more recent years, eDNA analysis has entered a new phase that of detecting not only the presence of organisms but also their behaviours. Particularly, there are high expectations for efficient and noninvasive detection of spawning activity through eDNA analysis because reproduction is one of the most important behaviours for maintaining species and populations in the life cycle of organisms^12–15^.

Many previous studies have reported that eDNA concentration increases substantially during the spawning season of fish and amphibians^e.g. 12,16–19^. Tsuji et al. (2021)^14^ demonstrated that significant diurnal variation in eDNA concentration is mainly caused by the released sperm during spawning events and it was used as evidence of spawning in a field survey targeting two Japanese medaka species (*Oryzias latipes* and *O. sakaizumii*). Furthermore, Bylemans et al. (2017)^13^ reported an increase in the ratio of nuclear DNA (nu-DNA) to mitochondrial DNA (mt-DNA) in collected eDNA when sperm was added to rearing water of Macquarie perch (*Macquaria australasica*). On the basis of these findings, attempts have been made to estimate the presence or absence of spawning activity and/or their scale by observing a spike in eDNA concentration or a change in the ratio before and after the expected spawning time^13,14,20,21^. It has been suggested that both approaches are useful and contribute to efficient and noninvasive detection of spawning; nevertheless, no studies have been conducted on the characteristics and dynamics of sperm-derived eDNA, which is key in both approaches.

The characteristics and dynamics of eDNA affect its detectability and persistence^22^. As eDNA analysis detects DNA fragments as a proxy for the presence of organisms, researchers have emphasised the importance of collecting the basic eDNA information to correctly interpret the results obtained by eDNA analysis^6,23,24^. Therefore, the knowledge of the origin, state, transport and fate of somatic-derived eDNA (e.g. skin, faeces, and mucus) has accumulated through numerous previous studies^6^. Nevertheless, the morphological features of sperm-derived eDNA are critically different from those of other somatic-derived eDNA examined in previous studies. For example, teleost spermatozoa range between 25 and 100 μm in length, and each consists of a head (approximately 2 μm), neck, neck section, middle section and tail^25^ (Fig. S1). The shape of the nucleus within the spermatozoa head is highly variable among taxonomic groups. The sperm head is tightly overlayed with the plasma membrane, but there is only a thin cytoplasmic layer between the plasma membrane and the nucleus^26^. The mitochondria are contained in the middle section of the flagellum around the axoneme, and they form a mitochondrial capsule with multiple disulphide bridges^25^. These characteristics can cause sperm-derived and somatic-derived eDNA to have different transports and fates.

This study focused on the particle size distribution (PSD) and persistence of sperm-derived eDNA. The size of eDNA is one of the factors that critically affect its dispersion distance and settling rate in water^27^, and it represents important information for understanding eDNA transport. Turner et al. (2014)^27^ demonstrated that the somatic-derived eDNA of common carp was most abundant in fractions with size between 1 μm and 10 μm. However, as the total length of teleost spermatozoa is much larger than 10 μm (it ranges from 25 μm to 100 μm), the abundance distribution of eDNA would be expected to shift its peak to larger fractions. Moreover, as the sperm nucleus is only overlayed with the plasma membrane and a thin cytoplasmic layer, it is likely to be degraded more rapidly by microbes and extracellular enzymes, being supplied to the smaller fractions. Additionally, if these speculations are correct, then the sperm-derived eDNA may have short-lived persistence in larger size fractions, which may allow for the selective collection of sperm derived-eDNA based only on size fraction for some time after spawning^28^. Time-limited and/or selective collection of sperm-derived DNA would help to detect spawning activity with higher sensitivity on the basis of the observation of spikes in concentration or ratio caused by sperm release. Furthermore, understanding the PSD of sperm-derived eDNA in relation to its persistence will provide completely new knowledge specific to the spawning season based on the characteristics and dynamics of eDNA.

The objective of this study was to examine the persistence and PSD of sperm-derived eDNA. Two experiments were conducted using the Japanese jack mackerel (*Trachurus japonicus*) as a model species. First, in experiment 1 (Exp. 1), the PSD and time-dependent degradation process of sperm-derived eDNA were investigated by artificially adding sperm to seawater, which was free from somatic-derived eDNA. Then, in experiment 2 (Exp. 2), Japanese jack mackerel were kept in tanks and the changes in DNA concentration and SPD before and after spawning activity were examined.

## Materials and Methods

Fig.1 shows the overview of the two experimental designs. Both experiments were performed at the Maizuru Fisheries Research Station (MFRS) of Kyoto University, Japan, located in front of Maizuru Bay.

**Fig. 1.**
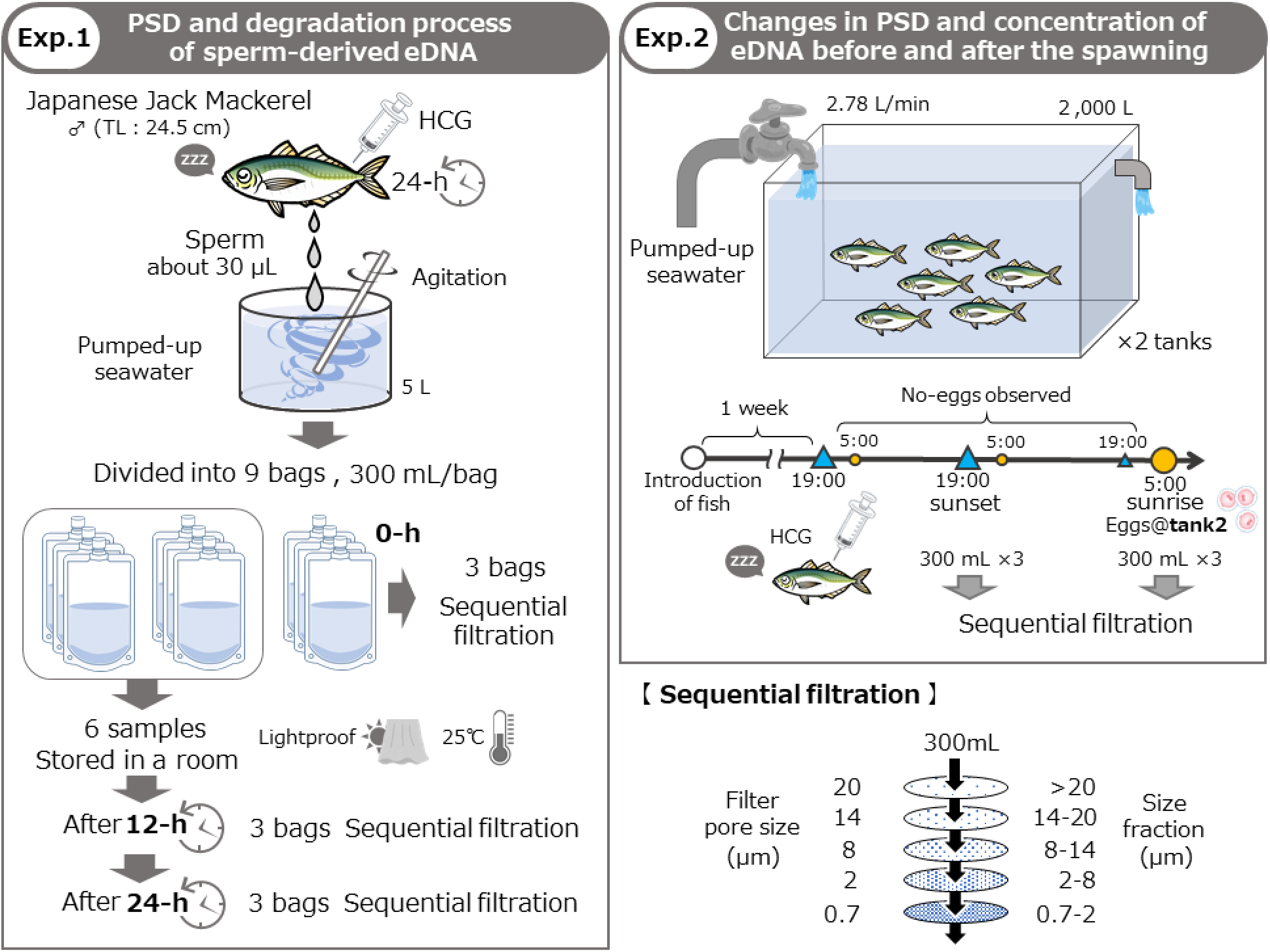
Overview of the two experimental designs adopted in this study.

### Experimentsl species and ethical statement

Japanese jack mackerel (*Trachurus japonicus*) was used as a model species in this study. It is one of the most commercially important fish in East Asia, including Japan^29^, and numerically it is the most dominant in Maizuru Bay from spring to autumn^30–32^. The main spawning grounds of Japanese jack mackerel are located in the East China Sea, but small to medium-scale spawning grounds are also scattered along Japanese coast^33,34^. Just outside Maizuru Bay, larvae as well as mature parental fish are generally captured^35^, suggesting that the species also spawn in and around the bay. Nonetheless, the timing and location of spawning have not been identified, and future surveys are desired from the perspective of resource management. The fish used for the experiments were captured in a commercial set net in Tai, Maizuru, on 29 May, 2020 and were transferred to MFRS. The sex of Japanese jack mackerel cannot be determined morphologically; therefore, after all the experiments were completed, the fish were anaesthetised and dissected, and their sex was determined through gonad observation. The current laws and guidelines in Japan regarding animal experiments allow the usage of fish without any ethical approval from any authority. However, all experiments were performed with maximal attention to animal welfare.

### Experiment 1: Particle size distribution and time-dependent degradation of sperm-derived eDNA

A Japanese jack mackerel (male, total length 24.5 cm) was injected with 83 IU of human chorionic gonadotropin (HCG; ASKA Pharmaceutical Co., Ltd., Tokyo, Japan) (Yoda et al. 2006). During the injection procedure, fish were anaesthetised with 2-phenoxyethanol (Hayashi Pure Chemical Ind., Ltd., Osaka, Japan). Twenty-four hours later, fish was re-anaesthetised and its abdomen was compressed to collect approximately 30 μL of sperm. The collected sperm was spiked in 5 L of fine-filtered seawater, which was pumped up from a depth of 6 m below the research station^36^. The seawater was agitated well and then dispensed into nine separate bags with caps (300 mL/bag; DP16-TN1000, Yanagi, Aichi, Japan). Of the nine seawater samples, three were selected as 0-h samples and were sequentially filtered through a series of filters with different pore sizes, i.e. 20, 14, 8, 2 μm (Track-Etched Membrane PCTE filter; GVS Japan, Tokyo, Japan) and 0.7 μm (GF/F glass-fibre filter; Cytiva, Tokyo, Japan). The remaining six seawater samples were stored in a room at 25°C, away from direct sunlight. After 12 and 24 h, in each time point, the stored three seawater samples were sequentially filtered as previously done for the 0-h samples. All the filters were immediately stored at −20 °C until DNA extraction.

### Experiment 2: Examination of the changes in PSD and eDNA concentration before and after spawning

Two tanks containing 2,000 L of fine-filtered pumped-up seawater were prepared. Six or seven Japanese jack mackerels were placed in each tank for about 1 week before Exp. 2 for them to acclimatise. The experiment was started on 5 June, 2020. Fish were anaesthetised with 2-phenoxyethanol and HCG was injected in appropriate quantities on the basis of the individual estimated body weight, which was calculated from the total length of each fish (Table S1). After HCG injection, the fish were immediately returned to their tank. Throughout the experiment, fine-filtered pumped-up seawater was poured into each tank at a rate that would replace the water within 12 h (i.e. 2.78 L/min). Twenty-four hours after all the fish had been injected with HCG (i.e. on 6 June, 2020, at 7:00 p.m.); no spawning was observed yet in either tank. As pre-spawning samples, three replicates of 300 mL of rearing water were collected from each tank (at 7:00 p.m.; hereafter referred to as ‘sunset’). The collected rearing water samples were sequentially filtered in the same manner as described in Exp. 1. The tanks were checked for the presence of any eggs at sunrise and sunset because Japanese jack mackerels spawn at night. On 8 June, 2020, at 5:00 a.m., eggs were observed in tank 2. Three replicates of 300 mL of rearing water were collected from each tank as after-spawning samples (at 5:00 a.m.; hereafter referred to as ‘sunrise’). The collected samples were sequentially filtered as previously done for the pre-spawning samples. At each time point during filtration, 300 mL of inlet seawater for tanks and ultrapure water were filtered using GF/F filters (Cytiva, Tokyo, Japan) as inlet- and filtration-negative controls to monitor unexpected contamination. All filters were immediately stored at −20 °C until DNA extraction.

### DNA extraction from each filter sample

The total eDNA on each filter was extracted following the method described by Tsuji with minor modification. First, each filter sample was placed in the upper part of the spin column (EconoSpin, EP-31201; GeneDesign, Inc., Osaka, Japan) with its silica-gel membrane removed. After centrifugation for 1 min at 6,000 g, each filter sample was moved to the 2 mL tube at the lower part of the spin column. A total of 420 μL of a solution, composed of 200 μL ultrapure water, 200 μL Buffer AL, and 20 μL proteinase K, was placed on each filter, and the spin columns were incubated for 45 min at 56 °C. After incubating, each filter sample was re-placed on the upper part of the spin column and centrifuged for 1 min at 6,000 g. 100 μL of Tris-EDTA buffer (pH 8.0) was placed on the filter and incubated for 1 min at 26 °C (room temperature). After centrifugation for 1 min at 6,000 g, 600 μL of ethanol was added to collected liquid and mixed well by gent pipetting. The DNA mixture was transferred to a DNeasy mini spin column (Qiagen, Hilden, Germany), and DNA was purified following the manufacturer’s protocol. The DNA was finally eluted in 100 μL of Buffer AE.

### Quantitative real-time PCR (qPCR) assays

Japanese Jack Mackerel’s eDNA concentrations at each size fraction were quantified using two types of species-specific primer-probe sets which were designed to amplify mitochondrial cytochrome b genes (Cyt *b*) and nuclear internal transcribed spacer-1 regions (ITS1) of ribosomal RNA genes, respectively (Table S2). The qPCR was performed with the same conditions for each target region individually. Each TaqMan reaction was conducted in a total volume of 15 μL, containing 900 nM of each primer, 125 nM TaqMan probe, 0.075 μL AmpErase uracil N-glycosylase (Thermo Fisher Scientific), 7.5 μL 2 × TaqMan Environmental Master Mix 2.0 (Thermo Fisher Scientific), and 2 μL DNA template. A standard dilution series containing 3 × 10^1^ to 3 × 10^4^ copies of each target region were used for qPCR runs. As PCR negative controls, 2.0 μL of ultrapure water was added and analyzed instead of the DNA template. The qPCR thermal conditions were as follows: 2 min at 50 °C, 10 min at 95 °C, then 55 cycles of 15 s at 95 °C, and 60 s at 60 °C. The qPCR was performed in triplicate for all eDNA samples, standard dilution series, and the PCR-negative control using a StepOne-Plus Real-Time PCR system (Applied Biosystems, FosterCity, CA, USA). The R^2^ values for the standard lines of all qPCR for Cyt *b* and ITS1 region ranged from 0.995 to 0.999 (Table S3). No amplification was observed in any of the filtration-and PCR-negative controls. The target DNA was detected from inlet-seawater in experiment 2, but the DNA copy number was so small compared to the amount of DNA detected in the tanks (< 0.7%) that it was deemed not to affect the results.

### Statistical analysis

All analyses were conducted in R ver. 3.6.0^37^ and the significance level was set at 0.05 in all of them. The statistical method for each data analysis was chosen on the basis of data distribution, as assessed by the Shapiro–Wilk test. In Exp. 1, first, the total copy number (+1) of Cyt b or ITS1 including all size fractions were compared among time points using the Kruskal–Wallis test followed by the Conover’s test with Holm adjustment (PMCMRplus package; ver. 1.9.3). In the same way, the ratio of ITS1 to Cyt *b* concentrations was compared among the time points. Subsequently, for each time point and DNA region, the number of DNA copies was compared among size fractions using the Kruskal–Wallis test followed by the Conover’s test with Holm adjustment. In Exp. 2, first, the total copy number of Cyt *b* or ITS1 detected in all size fractions were compared between the two time points (sunset and sunrise) using the t-test and/or Wilcoxon rank-sum test. The t-test method (i.e. the Student’s t-test or Welch’s t-test) was chosen on the basis of the variance equality of data as assessed by the F-test. When the Wilcoxon rank-sum test did not show significant differences, an additional t-test was performed to examine the significance of the differences more robustly. In the same way, for each tank, the ratio of ITS1 to Cyt *b* concentrations was compared between the time points. Then, for each DNA region and size fraction, the number of DNA copies between the time points was compared using the F test followed by the t-test and/or Wilcoxon rank-sum test. Similarly, the ratio of ITS1 to Cyt *b* concentrations was also compared between the time points.

## Results and Discussion

### Time-dependent degradation process with a shift in the PSD of sperm-derived eDNA

The total number of DNA copies significantly decreased with time in both Cyt *b* and ITS1 (Conover’s test; 0–12 and 12–24-h, *P* < 0.05; 0-24-h, *P* < 0.001 in both DNA regions; Figure 2A, B). Particularly, during the first 12 h, 95.9% of Cyt *b* and 99.6% of ITS1 were degraded. These results are consistent with previous studies using somatic-derived eDNA which suggested that the degradation begins immediately after release and is faster in nu-eDNA than mt-eDNA^38–40^. Moreover, it is important to note that the dynamics of eDNA in aquatic environments is affected by transport including settling, advection and dispersion as well as degradation^6^. Our laboratory experiment focused only on the degradation of sperm-derived eDNA. Hence, it is expected that the sperm-derived eDNA of Japanese jack mackerel released in a real aquatic environment will be almost undetectable in less than 12 h. However, as the degradation of sperm-derived eDNA was investigated for the first time in this study, there is no knowledge of the effects of species and physical parameters (e.g. temperature, pH, DO, salinity) on the degradation rate. Further insights into these aspects in future studies would allow a better understanding of the dynamics of sperm-derived eDNA.

**Fig. 2.**
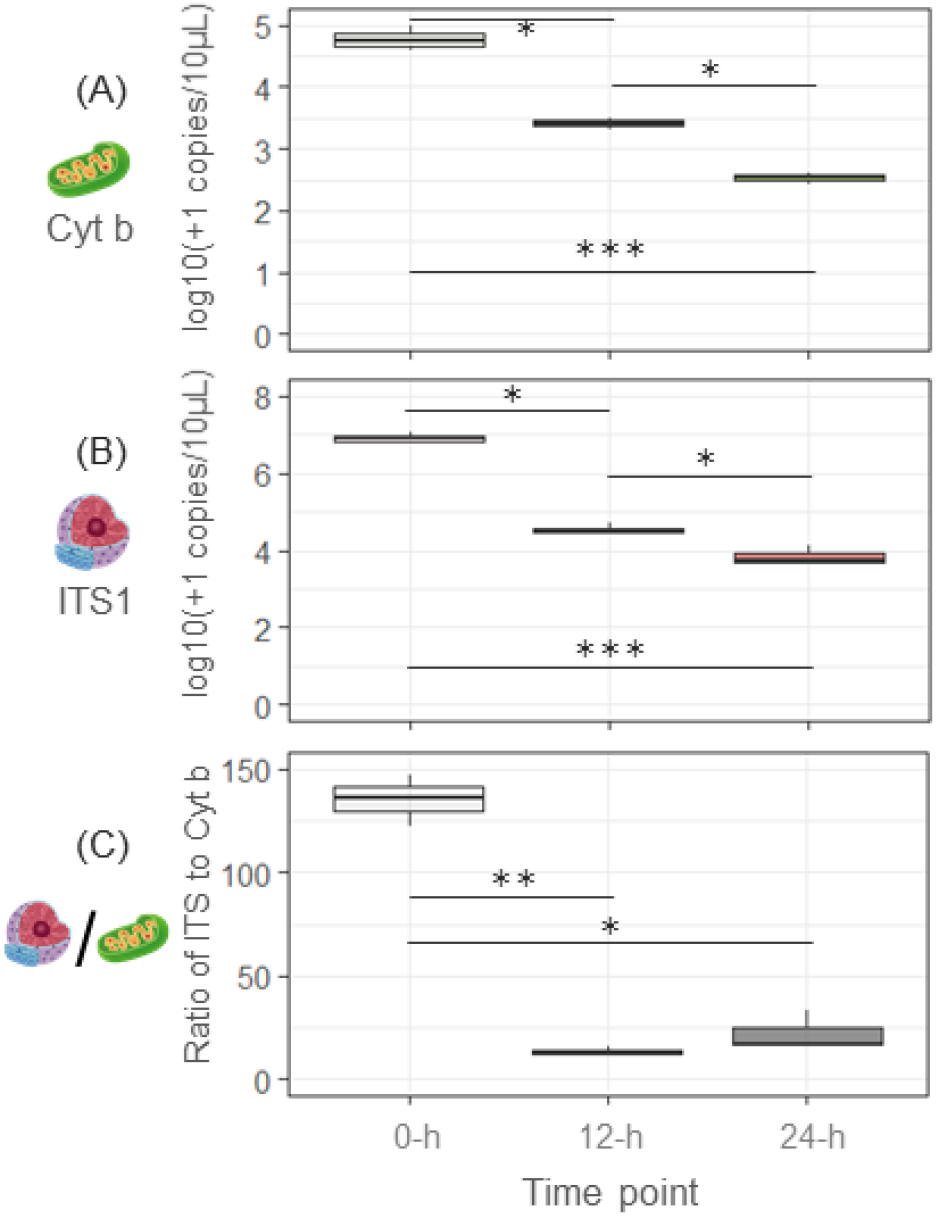
Temporal changes in the concentration of Japanese jack mackerel sperm-derived eDNA: (A) mt-eDNA, Cyt *b*, (B) nu-eDNA, ITS1 and (C) the ratio ITS1 to Cyt *b*. The DNA concentration is the sum of the copy numbers detected on the five filters with different pore sizes. Significant differences are indicated by asterisks (* *P* < 0.05, ** *P* < 0.01, *** *P* < 0.001; Conover’s test).

The ratio of ITS1 to Cyt *b* concentrations at 0-h was significantly greater than that at 12− and 24− h (Conover’s test; 0− and 12-h, *P* < 0.01; 0− and 24-h, *P* < 0.05; Fig. 2C), indicating that the concentration of nu-DNA (ITS1) in Japanese jack mackerel sperm was approximately 135-fold higher than that of mt-DNA (Cyt *b*). Considering that, when water contained only somatic-derived eDNA, the observed ITS1 to Cyt *b* ratio ranged from 5 to 12 (see Exp.2; Figure 4C, sunset), a 135-fold increase represents a very large ratio. This result was because the sperm contains less mt-DNA and more highly condensed nu-DNA^13^. Additionally, although the decrease in the ratio with time is reasonable (considering that the degradation rate of nu-eDNA is faster than that of mt-eDNA), it is worth noting that the ratio decreased rapidly to approximately 13.5 within 12 h (Fig. 2C). If only the changes in the nu-DNA to mt-DNA ratio are used as an indicator, it may be possible to detect spawning activity only for a few hours after the spawning time. These observations suggest that spawning surveys using ratio changes as an indicator should be carried out at the known spawning time of target species; otherwise, there is a high risk of falsenegative results. By contrast, a time-limited increase in the ratio may be useful for the detailed identification of the spawning timing and/or for monitoring the amount of spawn during very short periods of time.

As the total sperm length in Carangidae is approximately 44–49 μm, sperm-derived eDNA was expected to be recovered in larger size fractions. Nevertheless, the highest DNA concentrations were observed in the 0.7–2 μm size fraction in both DNA regions at all time points (Fig. 3A, B). Additionally, although the *P*-values obtained from Conover’s test were different, the frequency distribution of DNA concentrations detected in the other size fractions was also very similar between DNA regions at all time points (Fig. 3A, B; Table S4). These results should be related to the fact that sperm cells are mainly composed of an oval capsule-shaped head containing DNA and a long thin tail. The sperm head portion in Carangidae contains an elongated nucleus (long side, approximately 2 μm; short side, 1.2 μm) and some mitochondria (diameter, approximately 0.7 μm)^41^. Thus, the long thin tail did not inhibit the passage of the sperm through the filter, suggesting that the sperm could easily pass through a pore size larger than their head size. Based on this, the PSD of sperm-derived eDNA is expected to vary depending on the size of the sperm head. In the future, clarifying the relationship between sperm head size and/or shape and the PSD of sperm-derived eDNA for more species and taxa will contribute to a better understanding of the dynamics of sperm-derived eDNA.

**Fig. 3.**
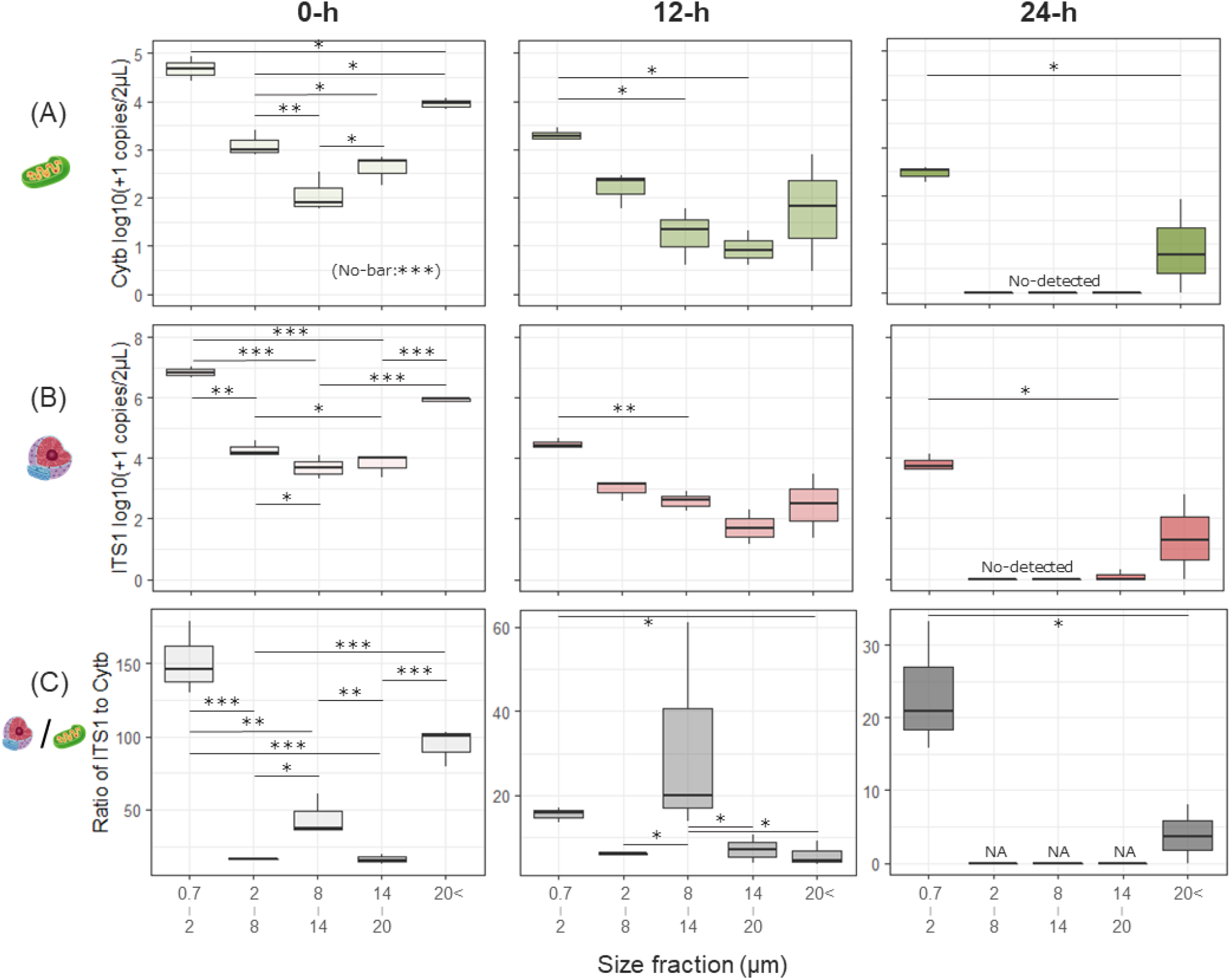
Temporal changes in the sperm-derived eDNA PSDs of Japanese jack mackerel: (A) mt-eDNA, Cyt *b*, (B) nu-eDNA, ITS1 and (C) the ratio ITS1 to Cyt *b*. Significant differences are indicated by asterisks (* *P* < 0.05, ** *P* < 0.01, *** *P* < 0.001; Conover’s test).

In both DNA regions, the concentration of sperm-derived eDNA decreased over time for all size fractions, whereas the PSD shifted from larger to smaller particle sizes over time (Fig. 3A and B, Fig. S2). This particle size shift has also been observed in a previous study that targeted somatic-derived eDNA of Japanese jack mackerel^22^. At the 0-h time point, the sperm head including nucleus and mitochondria should still be tightly overlayed with the plasma membrane, in all size fractions. However, over time, the plasma membrane will be lysed and fragmented because of the activity of microbes and exonucleases^22,42,43^. As sperm cell have a much thinner cytoplasmic layer than somatic cells^25^, the loss of the plasma membrane would easily release the nucleus and mitochondria into the environment. In this context, it is interesting to note that almost no DNA was detected for size fractions ranging between 2 μm and 20 μm at the 24-h time point. This indicates that the nucleus and mitochondria released into the environment may be rapidly degraded to a smaller size (< 2 μm) under the direct influence of enzymatic activity.

Regarding the ratio of ITS1 to Cyt *b* concentrations, if the nuclei and mitochondria in the sperm heads were trapped in each size fraction without separation, there should be no difference in the ratio among the size fractions; however, in fact, significant differences in the ratio were observed among almost all size fractions at 0-h (Fig. 3C, 0-h). This result indicates that the degradation of sperm-derived DNA, like that of somatic-derived eDNA^22^, begins with the release of the nucleus and mitochondria into the environment through the loss of the plasma membrane. Moreover, the observed PSD and ratio suggest that these released cell compornents have a size distribution that is not simply dependent on their individual sizes. As the size of mitochondria in sperm is approximately 0.7 μm, the released organelles were expected to be collected in the smallest size fraction, i.e. 0.7–2 μm. Nevertheless, the ratio observed in the >2 μm size fraction was much smaller than the overall ratio at 0 h (approximately 135; Fig. 2C, 0-h), suggesting that mitochondria were more likely to be trapped in larger size fractions than the nucleus (Fig. S2). This suggests that some of the mitochondria in the sperm would be separated from the nucleus in a state of interlocked each other ^cf.25^(Fig. S1). Additionally, a characteristic increase in the ratio was observed in the 8–14 μm size fraction up to the 12-h time point (Fig. 3C). However, as the sperm nuclei are approximately 2 μm in length (long side), it was unlikely that they could be recovered individually on an 8 μm pore size filter. In view of this, the observed increase in the ratio within the 8–14 μm size fraction may suggest that the nuclei were collected in a state where they were aggregated together for some reason. Further studies are needed to clarify the physical forms of sperm-derived eDNA released into natural environments in relation to degradation and PSD.

### Changes in DNA concentration and PSD before and after spawning activity

Only in tank 2, where fish spawned at night, significantly higher DNA concentrations were observed at sunrise than at sunset for both DNA regions (Cyt *b*, *P* < 0.01; ITS1, *P* < 0.05; Fig. 4A and B, Table S5). All the eggs collected from tank 2 at sunrise were fertilised, so there is no doubt that males released sperm during spawning events. These results confirm the findings of previous studies indicating that a spike in eDNA concentration before and after spawning can be used as evidence of spawning events^14,20,21^. Conversely, a significant decrease in the ratio between sunset and sunrise was observed in both tanks, even though an increase due to the presence of sperm-derived eDNA was expected in tank 2 (tank 1, *P* < 0.01; tank 2, *P* < 0.05; Fig. 4C, Table S5). This reduction in the ratio can be attributed to the following two facts: (1) the increased opportunities for physical contact among individuals due to the higher rearing density in the limited space and (2) the differences in degradation rate between nu-DNA and mt-DNA. First, as almost all fish in Exp. 2 had reached full maturity due to the HCG injection, the males in both tanks might have exhibited courtship behaviour towards females and/or aggressive behaviour towards other males, which involved physical contact during the night. Fish excitation and physical contact increase the number of epidermal cells and mucus, which are the main sources of eDNA, and in consequence, the somatic-derived eDNA concentration will temporarily increase^cf.22^. This assumption is supported by the fact that even tank 1, where no spawning was observed, also showed an upward trend in the concentration of both DNA regions (trending to significance; Cyt *b*, *P* = 0.10; ITS1, *P* = 0.36; Fig. 4A and B). Second, as nu-DNA released into water decays more rapidly than mt-DNA^44^, it is expected that the nu-DNA to mt-DNA ration of DNA released in large quantities during the night is lower at sunrise than in normal conditions. Consequently, as a higher amount of somatic-derived eDNA was temporarily released during the night, the nu-DNA to mt-DNA ratio observed at sunrise may be lower. For these reasons, the results obtained in this study question the usefulness of detecting real-spawning events, including courtship and threatening behaviour, based on the changes in the nu-DNA and mt-DNA ratio. However, it remains possible that the decrease in the ratio is only observed in laboratory settings (i.e. in tanks), and thus, this aspect remains to be examined in further studies conducted under natural conditions.

**Fig. 4.**
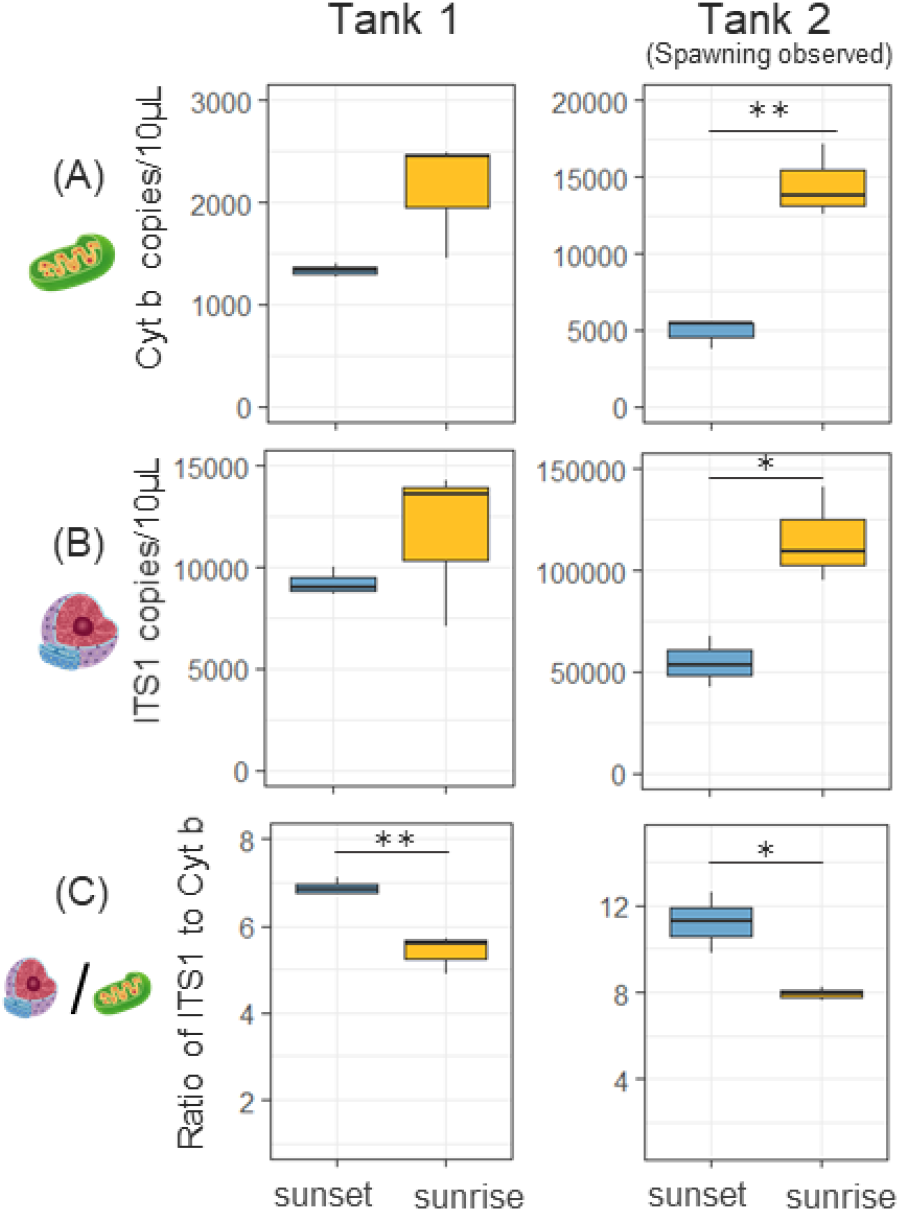
Comparisons of eDNA concentrations between sunset and sunrise in Exp. 2. The blue and orange box plots indicate sunset and sunrise, respectively. Only in tank 2, the fish spawned during the night. The DNA concentration is the sum of the copy numbers detected on the five filters with different pore sizes. Significant differences are indicated by asterisks (* *P* < 0.05, ** *P* < 0.01, *** *P* < 0.001; t-test or Wilcoxon rank-sum test).

**Fig. 5.**
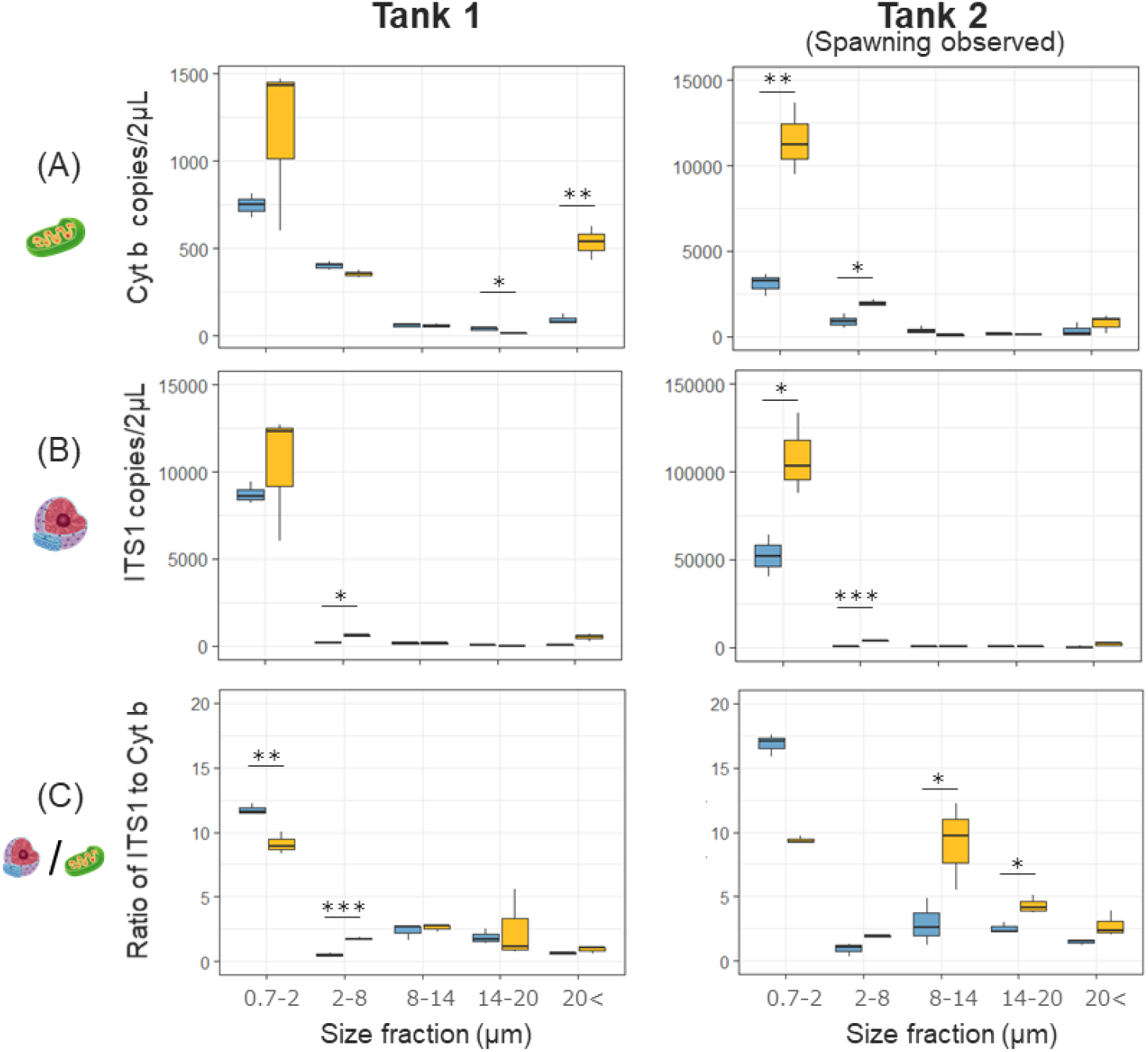
Comparisons of eDNA PSDs between sunset and sunrise in Exp. 2. The blue and orange box plots indicate sunset and sunrise, respectively. Only in tank 2, the fish spawned during the night. Significant differences are indicated by asterisks (* *P* < 0.05, ** *P* < 0.01, *** *P* < 0.001; t-test or Wilcoxon rank-sum test). It was not possible to determine whether the differences in DNA ratio in the 0.7–2 μm size fraction of tank 2 were significant (Wilcoxon rank-sum test, *P* = 0.1; additional t-test, *P* < 0.001).

The results of this study indicate that spawning activity shifts the PSD of eDNA into smaller size fractions, which would contribute to elucidating the characteristics and dynamics of eDNA specifically during the spawning season. For PSD at sunset and sunrise, the 0.7–2 μm size fraction always had the highest concentration, regardless of the DNA region and whether spawning was observed or not. Nevertheless, a significant increase in the concentration of both DNA regions was observed in the 0.7–2 and 2–8 μm size fractions only in tank 2, possibly due to the effect of the sperm released during spawning activity. These results are consistent with the fact that the PSD of sperm-derived DNA was always most abundant in the 0.7–2 μm size fraction for both DNA regions (Exp. 1, Fig. 3). Conversely, based on the observations in Exp. 1 using sperm-derived eDNA, an increase in the concentrations of both DNA regions was expected in the size fractions 14 ≤ μm in tank 2 at sunrise, but this did not occur. In Exp 2, as it was difficult to monitor the spawning directly, it is possible that the sperm-derived eDNA was recovered in an advanced state of degradation. Additionally, because the rearing water in tanks was constantly being replaced (2.78 L/min), the released sperm-derived eDNA would have been diluted. For these reasons, it is believed that sperm-derived eDNA with shorter persistence was not detected from larger size fractions as much as it had been predicted. Interestingly, a characteristic increase in the ratio of the 8–14 μm size fraction after spawning, which was observed in Exp. 1 using sperm-derived eDNA, was also observed in tank 2. Additionally, only in tank 2, an increase in the ratio of the 14–20 μm size fraction was also observed after spawning. Nevertheless, the concentrations of both DNA regions in the 8–14 μm and 14–20 μm size fractions did not significantly increase between before and after the spawning activity; instead, the concentration of mitochondria showed a slight downward trend. For these reasons, although the presence of sperm-derived eDNA may alter the ratio of nuclei to mitochondria in certain size fractions, the detailed mechanisms and validity of the results must to be further investigated.

### Implications and Perspectives

It was observed that the degradation of sperm-derived eDNA proceeded rapidly with a shift to smaller particle sizes, regardless of the DNA region (Cyt *b* and ITS1), indicating that spawning activity temporarily shifts the overall PSD of eDNA into a smaller size. Furthermore, the observed PSD and ratio suggested that the released nuclei and mitochondria from sperm show a size distribution that is not simply dependent on their individual sizes. The present study is the first report to investigate the persistence and PSD of sperm-derived eDNA, as well as the first to reveal the characteristics and dynamics of sperm-derived eDNA. The results obtained would contribute to the elucidation of both, specifically during the spawning season, and can be the basis for the further development of eDNA-based monitoring of spawning.

It was here confirmed that the spike in eDNA concentration and change in the nu-DNA to mt-DNA ratio (which have been used as indicators in previous eDNA analysis based studies) are definitely caused by spawning activity. On the basis of the findings obtained on the characteristics and dynamics of sperm-derived eDNA, various considerations can be made for spawning surveys that rely on the observation of spikes in eDNA concentrations and ratio changes, which have been established in previous studies^13,14,21^. First, when detecting spikes in concentration caused by sperm-derived eDNA, it is necessary to use a filter with a pore size smaller than the short side of the nucleus and mitochondria within the sperm head of the target species. The selection of a filter with an appropriate pore size allows us to reliably trap sperm-derived eDNA and robustly detect spikes in DNA concentration. Then, when changes in nu-DNA to mt-DNA ratio are used as an indicator of spawning activity, it is necessary to be aware of the uncertainty and persistence of these ratio changes. The results of this study showed that the changes may only be observed in the first few hours after spawning (within less than 12 h). Additionally, changes in the ratio can easily be caused by differences in the capture rate of each organelle depending on the filter pore size, and by the excessive release of somatic-derived eDNA due to physical contact among individuals, including the parents. The usefulness and certainty of this approach are open to further investigation.

Although some aspects of the characteristics and dynamics of sperm-derived eDNA were elucidated, there remain knowledge gaps that must be addressed to perform more accurate and robust spawning surveys in natural environments. For example, our results suggest that sperm-derived eDNA is degraded more rapidly than somatic-derived eDNA; nevertheless, the degradation rate, its relationship to environmental factors (such as water temperature) and differences between species need to be investigated. Additionally, the morphology, size and number of organelles (i.e. nuclei and mitochondria) of sperm heads vary widely among species and taxa (ex. fish^45^, length 1.2–5.7 μm, width 0.6–1.9 μm; frog^46^, length 10.7–26.8 μm, width 1.0–2.0 μm). These differences will have a significant impact on the persistence and PSD of sperm-derived DNA. The understanding and accumulation of basic information on the characteristics and dynamics of sperm-derived eDNA will maximise the applicability and usefulness of eDNA analysis to spawning surveys. A novel, efficient and noninvasive strategy for detecting spawning activity using eDNA analysis would represent a powerful tool for the conservation and management of ecosystems.

## Authors’ contributions

S.T. conceived and designed research. All authors performed tank experiments including fish rearing and water sampling. S.T. performed molecular experiments and data analysis. S.T. wrote the early draft and completed it with significant inputs from H.M. and R.M.

## Data availability

Full details of the qPCR results for each experiment of the present study are available in the supporting information (Table S6 and S7).

## Acknowledgements

We thank Ms Masako Hara for her valuable advice. This study was supported by JSPS KAKENHI Grant Number JP20K15578. We declare no conflicts of interests.

---

**Fig. S1.**
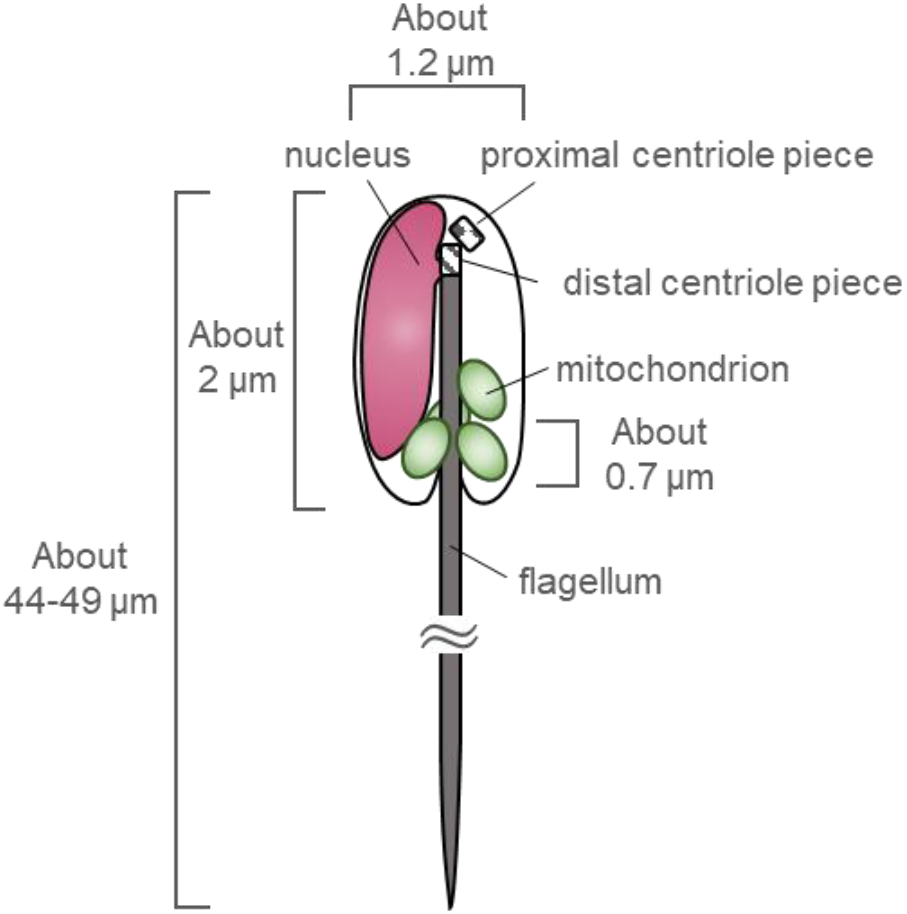
Simple schematic representation of the ultrastructure of Carangidae sperm (modified from Ulloa-Rodríguez et al. 2017).

**Fig. S2.**
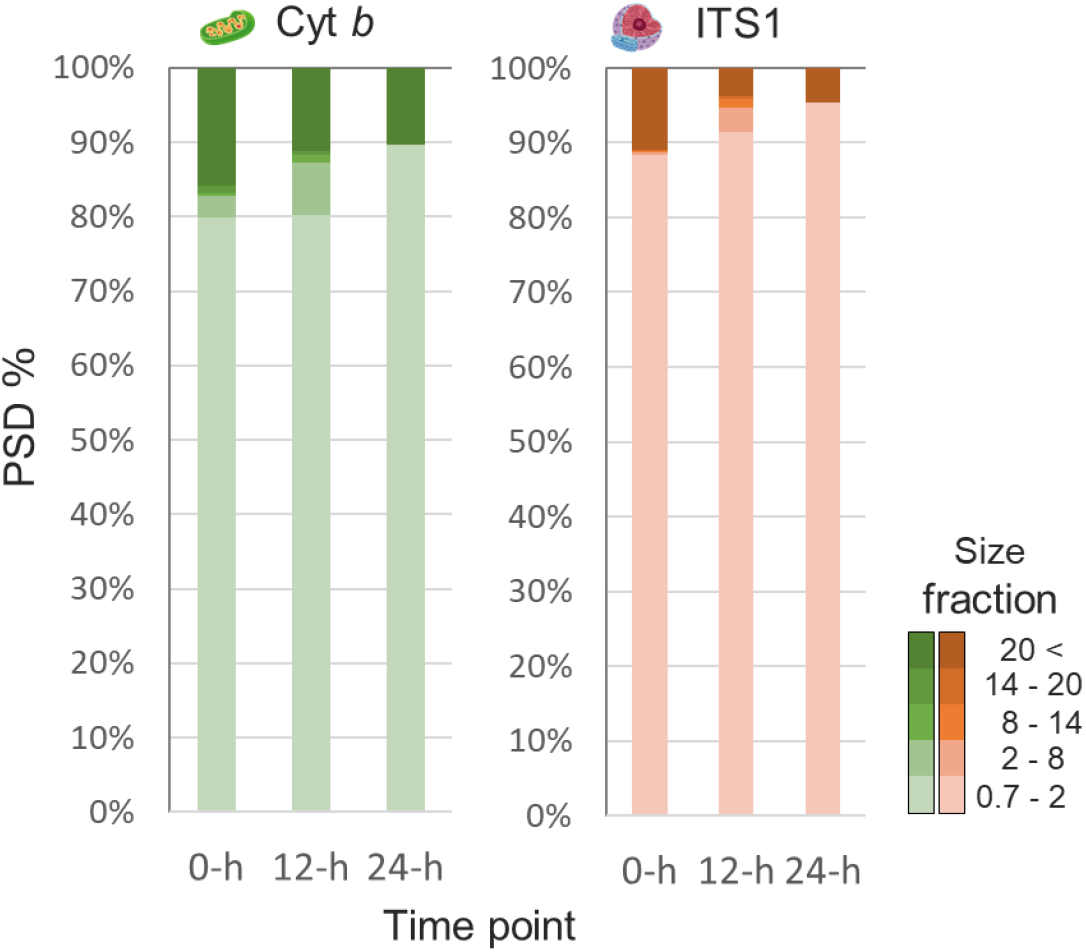
The temporal changes in percentage of copy number in each size fraction relative to the total number of copies of sperm-derived eDNA.

