## Supplemental Tables for "Analysis of the persistence and particle size distributional shift of sperm-derived environmental DNA to monitor Jack Mackerel spawning activity"

Table S1 Detailed information on experimental fish and the injected amount of HCG. GSI is gonadal weight as apercentage of total body weight.

| Tank ID | Fish ID | sex | Total Length (cm) | Total body weight (g) | Gonadal weight (g) | GSI | Injected amount of HCG (IU) |
|---|---|---|---|---|---|---|---|
| Tank 1 | 1 | female | 28.7 | 258.1 | 5.5 | 2.1 | 146 |
| | 2 | female | 38.2 | 687.2 | 144.2 | 21.0 | 329 |
| | 3 | male | 28.6 | 284.8 | 18.7 | 6.6 | 146 |
| | 4 | male | 34.4 | 364.2 | 10.2 | 2.8 | 236 |
| | 5 | male | 28.2 | 257.3 | 9.7 | 3.8 | 132 |
| | 6 | male | 28.0 | 216.5 | 3.2 | 1.5 | 132 |
| | 7 | male | 33.7 | 336.4 | 4.8 | 1.4 | 236 |
| Tank 2 | 1 | male | 28.7 | 247.8 | 8.9 | 3.6 | 146 |
| | 2 | female | 29.3 | 291.5 | 10.3 | 3.5 | 146 |
| | 3 | male | 33.0 | 358.5 | 8.9 | 2.5 | 216 |
| | 4 | male | 32.5 | 330.3 | 10.5 | 3.2 | 216 |
| | 5 | male | 29.6 | 279.3 | 11.8 | 4.2 | 162 |
| | 6 | female | 35.7 | 463.5 | 27.6 | 6.0 | 280 |

Table S2 Detailed information on Japanese Jack Mackerel’s species specific primers/probe sets used in this study.

| Primers/probe | Target region | Sequences (5' −> 3') | Tm (°C) | Amplicon length (bp) | References |
|---|---|---|---|---|---|
| Tja_Cytb_F | | CAG ATA TCG CAA CCG CCT TT | 58.7 | | |
| Tja_Cytb_R | Cyt b (mt-DNA) | CCG ATG TGA AGG TAA ATG CAA A | 59.8 | 164 | Minamoto et al. (unpublished), Yamamoto et al. (2016) |
| Tja_Cytb_Pr | | [FAM]-TAT GCA CGC CAA CGG CGC CT-[BHQ1] | 67.9 | | |
| Tja_ITS1_F | | GCG GGT ACC CAA CTC TCT TC | 60.1 | | |
| Tja_ITS1_R | ITS1 (nu-DNA) | CCT GAG CGG CAC ATG AGA G | 63.2 | 164 | Jo et al. (2019) |
| Tja_ITS1_Pr | | [FAM]-CTC TCG CTT CTC CGA CCC CGG TCG-[BHQ1] | 70.8 | | |

Table S3 Detailed information on qPCR qualities. qPCR_ID corresponds to that at Table S6 and S7.

| Exp.No | qPCR_ID | Slope | Y-inter | R^2 | Eff (%) |
|---|---|---|---|---|---|
| 1 | PCR01 | -3.43 | 40.72 | 0.996 | 95.60 |
| 1 | PCR02 | -3.48 | 40.91 | 0.999 | 93.96 |
| 1 | PCR03 | -3.82 | 43.78 | 0.998 | 82.82 |
| 1 | PCR04 | -3.82 | 44.10 | 0.996 | 82.61 |
| 1 | PCR05 | -3.84 | 44.38 | 0.997 | 82.06 |
| 2 | PCR01 | -3.26 | 39.86 | 0.999 | 102.64 |
| 2 | PCR02 | -3.82 | 43.78 | 0.998 | 82.82 |
| 2 | PCR03 | -3.34 | 40.47 | 0.998 | 99.20 |
| 2 | PCR04 | -3.87 | 44.67 | 0.999 | 81.41 |
| 2 | PCR05 | -3.62 | 41.40 | 0.997 | 88.89 |
| 2 | PCR06 | -3.60 | 41.21 | 0.999 | 89.52 |
| 2 | PCR07 | -4.51 | 45.24 | 0.997 | 76.34 |
| 2 | PCR08 | -3.93 | 43.93 | 0.997 | 79.67 |
| 2 | PCR09 | -3.77 | 41.83 | 0.995 | 84.05 |

Table S4 Results of Shapiro-Wilk test and Conover's tests in Exp. 1, where bold values represent the statistical significances ($p < 0.05$).

| | | 0-h | | | | | 12-h | | | | | 24-h | | | | |
|---|---|---|---|---|---|---|---|---|---|---|---|---|---|---|---|---|
| | | Shapiro-Wilk test | 0.7 µm | 2 µm | 8 µm | 14 µm | Shapiro-Wilk test | 0.7 µm | 2 µm | 8 µm | 14 µm | Shapiro-Wilk test | 0.7 µm | 2 µm | 8 µm | 14 µm |
| Cyt b | Shapiro-Wilk test | **0.00003** | | | | | **0.0001** | | | | | **0.00002** | | | | |
| | 2 µm | | **0.001** | - | - | - | | 0.534 | - | - | - | | **0.001** | - | - | - |
| | 8 µm | | **0.000** | **0.001** | - | - | | **0.037** | 0.534 | - | - | | **0.001** | 1.000 | - | - |
| | 14 µm | | **0.000** | **0.032** | **0.044** | - | | **0.017** | 0.244 | 1.000 | - | | **0.001** | 1.000 | 1.000 | - |
| | 20 µm | | **0.042** | **0.042** | **0.000** | **0.001** | | 0.153 | 1.000 | 1.000 | 0.695 | | **0.049** | 0.079 | 0.079 | 0.079 |
| ITS1 | Shapiro-Wilk test | **0.00002** | | | | | **0.00001** | | | | | **0.000004** | | | | |
| | 2 µm | | **0.004** | - | - | - | | 0.455 | - | - | - | | **0.005** | - | - | - |
| | 8 µm | | **0.000** | **0.019** | - | - | | 0.112 | 0.853 | - | - | | **0.005** | 1.000 | - | - |
| | 14 µm | | **0.000** | **0.024** | 0.795 | - | | **0.009** | 0.160 | 0.603 | - | | **0.016** | 1.000 | 1.000 | - |
| | 20 µm | | 0.111 | 0.111 | **0.001** | **0.001** | | 0.099 | 0.853 | 0.890 | 0.603 | | 0.177 | 0.209 | 0.209 | 0.657 |
| ITS1/Cyt b | Shapiro-Wilk test | **0.021** | | | | | **0.00004** | | | | | **0.00005** | | | | |
| | 2 µm | | **0.000** | - | - | - | | 0.062 | - | - | - | | **0.001** | - | - | - |
| | 8 µm | | **0.003** | **0.032** | - | - | | 1.000 | **0.025** | - | - | | **0.001** | 1.000 | - | - |
| | 14 µm | | **0.000** | 0.428 | **0.010** | - | | 0.117 | 1.000 | **0.043** | - | | **0.001** | 1.000 | 1.000 | - |
| | 20 µm | | 0.098 | **0.001** | 0.098 | **0.000** | | **0.038** | 1.000 | **0.013** | 1.000 | | **0.049** | 0.079 | 0.079 | 0.079 |

Table S5 Results of Statistical analysis in Exp. 2, where bold values represent the statistical significances ($p < 0.05$).

| Region | Tank ID | test | Total copy number | 0.7 µm | 2 µm | 8 µm | 14 µm | 20 µm |
|---|---|---|---|---|---|---|---|---|
| Cyt b | Tank1 | Shapiro-Wilk test | **0.01** | 0.05 | 0.93 | 0.67 | 0.36 | 0.12 |
| | | F-test | - | **0.04** | 0.89 | 0.68 | **0.03** | 0.15 |
| | | t-test | - | 0.27 | 0.07 | 1.00 | **0.03** | **0.00** |
| | | Wilcoxon rank sum test | 0.10 | - | - | - | - | - |
| | Tank2 | Shapiro-Wilk test | 0.28 | 0.25 | 0.56 | 0.05 | 0.39 | 0.17 |
| | | F test | 0.28 | 0.19 | 0.27 | **0.02** | 0.75 | 0.71 |
| | | t-test | **0.00** | **0.00** | **0.01** | 0.18 | 0.87 | 0.33 |
| | | Wilcoxon rank sum test | - | - | - | - | - | - |
| ITS1 | Tank1 | Shapiro-Wilk test | 0.35 | 0.59 | 0.12 | 0.14 | 0.54 | 0.06 |
| | | F-test | 0.06 | **0.05** | 0.89 | **0.02** | 0.47 | **0.00** |
| | | t-test | 0.36 | 0.54 | **0.00** | 0.84 | 0.10 | 0.07 |
| | | Wilcoxon rank sum test | - | - | - | - | - | - |
| | Tank2 | Shapiro-Wilk test | 0.80 | 0.83 | 0.08 | 0.11 | 0.97 | 0.26 |
| | | F test | 0.45 | 0.43 | 0.74 | 0.36 | 0.66 | 0.64 |
| | | t-test | **0.02** | **0.02** | **0.00** | 0.78 | 0.32 | 0.15 |
| | | Wilcoxon rank sum test | - | - | - | - | - | - |
| ITS1/Cyt b ratio | Tank1 | Shapiro-Wilk test | | 0.41 | 0.06 | 0.09 | 0.05 | 0.21 |
| | | F-test | | 0.35 | 0.87 | 0.48 | 0.09 | 0.34 |
| | | t-test | | **0.01** | **0.00** | 0.52 | 0.72 | 0.17 |
| | | Wilcoxon rank sum test | - | - | - | - | - | - |
| | Tank2 | Shapiro-Wilk test | - | **0.04** | 0.23 | 0.66 | 0.63 | 0.15 |
| | | F test | - | - | **0.02** | 0.45 | 0.53 | 0.08 |
| | | t-test | - | - | 0.07 | **0.05** | **0.02** | 0.10 |
| | | Wilcoxon rank sum test | - | 0.10 | - | - | - | - |

Table S6 All raw data of qPCR (DNA copies per 2 μL template DNA) in Experiment 1.

| qPCR ID | Time point | Replication No. | Filter pore size | Region | $C_T$ | $C_T$ Mean | $C_T$ SD | Quantity | Quantity Mean | Quantity SD |
|---|---|---|---|---|---|---|---|---|---|---|
| PCR01 | STANDARD | | | Cyt b | 25.3 | 25.3 | 0.0 | 30000.0 | | |
| PCR01 | STANDARD | | | Cyt b | 25.3 | 25.3 | 0.0 | 30000.0 | | |
| PCR01 | STANDARD | | | Cyt b | 25.2 | 25.3 | 0.0 | 30000.0 | | |
| PCR01 | STANDARD | | | Cyt b | 28.8 | 28.8 | 0.0 | 3000.0 | | |
| PCR01 | STANDARD | | | Cyt b | 28.8 | 28.8 | 0.0 | 3000.0 | | |
| PCR01 | STANDARD | | | Cyt b | 28.8 | 28.8 | 0.0 | 3000.0 | | |
| PCR01 | STANDARD | | | Cyt b | 32.2 | 32.3 | 0.1 | 300.0 | | |
| PCR01 | STANDARD | | | Cyt b | 32.3 | 32.3 | 0.1 | 300.0 | | |
| PCR01 | STANDARD | | | Cyt b | 32.4 | 32.3 | 0.1 | 300.0 | | |
| PCR01 | STANDARD | | | Cyt b | 36.0 | 35.6 | 0.5 | 30.0 | | |
| PCR01 | STANDARD | | | Cyt b | 35.0 | 35.6 | 0.5 | 30.0 | | |
| PCR01 | STANDARD | | | Cyt b | 35.6 | 35.6 | 0.5 | 30.0 | | |
| PCR01 | 0 | 1 | 0.7 | Cyt b | 23.8 | 23.8 | 0.1 | 84412.5 | 84273.2 | 3197.6 |
| PCR01 | 0 | 1 | 0.7 | Cyt b | 23.8 | 23.8 | 0.1 | 87398.9 | 84273.2 | 3197.6 |
| PCR01 | 0 | 1 | 0.7 | Cyt b | 23.9 | 23.8 | 0.1 | 81008.3 | 84273.2 | 3197.6 |
| PCR01 | 0 | 1 | 2 | Cyt b | 30.4 | 30.5 | 0.1 | 1007.0 | 962.3 | 79.6 |
| PCR01 | 0 | 1 | 2 | Cyt b | 30.4 | 30.5 | 0.1 | 1009.5 | 962.3 | 79.6 |
| PCR01 | 0 | 1 | 2 | Cyt b | 30.6 | 30.5 | 0.1 | 870.4 | 962.3 | 79.6 |
| PCR01 | 0 | 1 | 8 | Cyt b | 34.3 | 34.2 | 0.1 | 73.3 | 77.3 | 3.7 |
| PCR01 | 0 | 1 | 8 | Cyt b | 34.2 | 34.2 | 0.1 | 80.5 | 77.3 | 3.7 |
| PCR01 | 0 | 1 | 8 | Cyt b | 34.2 | 34.2 | 0.1 | 78.0 | 77.3 | 3.7 |
| PCR01 | 0 | 1 | 14 | Cyt b | 32.9 | 33.0 | 0.1 | 188.1 | 179.3 | 16.3 |
| PCR01 | 0 | 1 | 14 | Cyt b | 32.9 | 33.0 | 0.1 | 189.3 | 179.3 | 16.3 |
| PCR01 | 0 | 1 | 14 | Cyt b | 33.2 | 33.0 | 0.1 | 160.4 | 179.3 | 16.3 |
| PCR01 | 0 | 1 | 20 | Cyt b | 26.8 | 26.8 | 0.0 | 11183.2 | 11558.5 | 379.1 |
| PCR01 | 0 | 1 | 20 | Cyt b | 26.7 | 26.8 | 0.0 | 11941.4 | 11558.5 | 379.1 |
| PCR01 | 0 | 1 | 20 | Cyt b | 26.8 | 26.8 | 0.0 | 11551.0 | 11558.5 | 379.1 |
| PCR01 | 0 | 2 | 0.7 | Cyt b | 24.6 | 24.7 | 0.0 | 48260.0 | 47411.0 | 739.7 |
| PCR01 | 0 | 2 | 0.7 | Cyt b | 24.7 | 24.7 | 0.0 | 46905.7 | 47411.0 | 739.7 |
| PCR01 | 0 | 2 | 0.7 | Cyt b | 24.7 | 24.7 | 0.0 | 47067.2 | 47411.0 | 739.7 |

| | | | | | | | | | | |
|---|---|---|---|---|---|---|---|---|---|---|
| PCR01 | 0 | 2 | 2 | Cyt b | 30.7 | 30.8 | 0.1 | 826.1 | 795.0 | 39.3 |
| PCR01 | 0 | 2 | 2 | Cyt b | 30.7 | 30.8 | 0.1 | 808.1 | 795.0 | 39.3 |
| PCR01 | 0 | 2 | 2 | Cyt b | 30.9 | 30.8 | 0.1 | 750.8 | 795.0 | 39.3 |
| PCR01 | 0 | 2 | 8 | Cyt b | 32.0 | 32.0 | 0.0 | 341.6 | 342.7 | 10.5 |
| PCR01 | 0 | 2 | 8 | Cyt b | 32.0 | 32.0 | 0.0 | 353.7 | 342.7 | 10.5 |
| PCR01 | 0 | 2 | 8 | Cyt b | 32.1 | 32.0 | 0.0 | 332.7 | 342.7 | 10.5 |
| PCR01 | 0 | 2 | 14 | Cyt b | 31.0 | 31.0 | 0.0 | 688.2 | 689.8 | 3.0 |
| PCR01 | 0 | 2 | 14 | Cyt b | 31.0 | 31.0 | 0.0 | 688.1 | 689.8 | 3.0 |
| PCR01 | 0 | 2 | 14 | Cyt b | 31.0 | 31.0 | 0.0 | 693.3 | 689.8 | 3.0 |
| PCR01 | 0 | 2 | 20 | Cyt b | 27.5 | 27.6 | 0.1 | 7127.5 | 6866.8 | 443.7 |
| PCR01 | 0 | 2 | 20 | Cyt b | 27.5 | 27.6 | 0.1 | 7118.3 | 6866.8 | 443.7 |
| PCR01 | 0 | 2 | 20 | Cyt b | 27.7 | 27.6 | 0.1 | 6354.5 | 6866.8 | 443.7 |
| PCR01 | 0 | 3 | 0.7 | Cyt b | 25.5 | 25.6 | 0.1 | 26757.8 | 26049.8 | 899.3 |
| PCR01 | 0 | 3 | 0.7 | Cyt b | 25.6 | 25.6 | 0.1 | 25037.9 | 26049.8 | 899.3 |
| PCR01 | 0 | 3 | 0.7 | Cyt b | 25.5 | 25.6 | 0.1 | 26353.7 | 26049.8 | 899.3 |
| PCR01 | 0 | 3 | 2 | Cyt b | 29.1 | 29.1 | 0.0 | 2489.5 | 2467.9 | 22.4 |
| PCR01 | 0 | 3 | 2 | Cyt b | 29.1 | 29.1 | 0.0 | 2469.5 | 2467.9 | 22.4 |
| PCR01 | 0 | 3 | 2 | Cyt b | 29.1 | 29.1 | 0.0 | 2444.7 | 2467.9 | 22.4 |
| PCR01 | 0 | 3 | 8 | Cyt b | 34.7 | 34.7 | 0.0 | 56.5 | 57.9 | 1.3 |
| PCR01 | 0 | 3 | 8 | Cyt b | 34.6 | 34.7 | 0.0 | 59.1 | 57.9 | 1.3 |
| PCR01 | 0 | 3 | 8 | Cyt b | 34.7 | 34.7 | 0.0 | 58.1 | 57.9 | 1.3 |
| PCR01 | 0 | 3 | 14 | Cyt b | 31.1 | 31.3 | 0.2 | 635.5 | 574.6 | 75.3 |
| PCR01 | 0 | 3 | 14 | Cyt b | 31.2 | 31.3 | 0.2 | 597.8 | 574.6 | 75.3 |
| PCR01 | 0 | 3 | 14 | Cyt b | 31.5 | 31.3 | 0.2 | 490.4 | 574.6 | 75.3 |
| PCR01 | 0 | 3 | 20 | Cyt b | 27.3 | 27.2 | 0.1 | 8313.3 | 8952.6 | 633.9 |
| PCR01 | 0 | 3 | 20 | Cyt b | 27.2 | 27.2 | 0.1 | 8963.6 | 8952.6 | 633.9 |
| PCR01 | 0 | 3 | 20 | Cyt b | 27.1 | 27.2 | 0.1 | 9581.0 | 8952.6 | 633.9 |
| PCR01 | 12 | 1 | 0.7 | Cyt b | 28.9 | 28.9 | 0.1 | 2756.8 | 2801.3 | 114.2 |
| PCR01 | 12 | 1 | 0.7 | Cyt b | 28.8 | 28.9 | 0.1 | 2931.0 | 2801.3 | 114.2 |
| PCR01 | 12 | 1 | 0.7 | Cyt b | 28.9 | 28.9 | 0.1 | 2716.0 | 2801.3 | 114.2 |
| PCR01 | 12 | 1 | 2 | Cyt b | 32.1 | 32.3 | 0.2 | 328.2 | 294.3 | 31.9 |
| PCR01 | 12 | 1 | 2 | Cyt b | 32.3 | 32.3 | 0.2 | 290.1 | 294.3 | 31.9 |
| PCR01 | 12 | 1 | 2 | Cyt b | 32.4 | 32.3 | 0.2 | 264.8 | 294.3 | 31.9 |

| | | | | | | | | | | |
|---|---|---|---|---|---|---|---|---|---|---|
| PCR01 | 12 | 1 | 8 | Cyt b | 35.0 | 34.7 | 0.4 | 45.3 | 56.6 | 14.4 |
| PCR01 | 12 | 1 | 8 | Cyt b | 34.8 | 34.7 | 0.4 | 51.8 | 56.6 | 14.4 |
| PCR01 | 12 | 1 | 8 | Cyt b | 34.3 | 34.7 | 0.4 | 72.8 | 56.6 | 14.4 |
| PCR01 | 12 | 1 | 14 | Cyt b | Undetermined | 38.9 | 0.1 | | | |
| PCR01 | 12 | 1 | 14 | Cyt b | 38.8 | 38.9 | 0.1 | 3.7 | 3.5 | 0.3 |
| PCR01 | 12 | 1 | 14 | Cyt b | 39.0 | 38.9 | 0.1 | 3.2 | 3.5 | 0.3 |
| PCR01 | 12 | 2 | 0.7 | Cyt b | 29.6 | 29.5 | 0.1 | 1741.0 | 1885.9 | 125.5 |
| PCR01 | 12 | 2 | 0.7 | Cyt b | 29.4 | 29.5 | 0.1 | 1959.9 | 1885.9 | 125.5 |
| PCR01 | 12 | 2 | 0.7 | Cyt b | 29.4 | 29.5 | 0.1 | 1956.9 | 1885.9 | 125.5 |
| PCR01 | 12 | 2 | 2 | Cyt b | 34.8 | 34.7 | 0.1 | 54.4 | 58.8 | 3.9 |
| PCR01 | 12 | 2 | 2 | Cyt b | 34.6 | 34.7 | 0.1 | 60.1 | 58.8 | 3.9 |
| PCR01 | 12 | 2 | 2 | Cyt b | 34.6 | 34.7 | 0.1 | 61.8 | 58.8 | 3.9 |
| PCR01 | 12 | 2 | 8 | Cyt b | 36.3 | 36.2 | 0.2 | 19.9 | 20.9 | 3.3 |
| PCR01 | 12 | 2 | 8 | Cyt b | 35.9 | 36.2 | 0.2 | 24.6 | 20.9 | 3.3 |
| PCR01 | 12 | 2 | 8 | Cyt b | 36.4 | 36.2 | 0.2 | 18.1 | 20.9 | 3.3 |
| PCR01 | 12 | 2 | 14 | Cyt b | 36.3 | 36.3 | 0.3 | 19.8 | 19.2 | 3.9 |
| PCR01 | 12 | 2 | 14 | Cyt b | 36.7 | 36.3 | 0.3 | 15.1 | 19.2 | 3.9 |
| PCR01 | 12 | 2 | 14 | Cyt b | 36.1 | 36.3 | 0.3 | 22.8 | 19.2 | 3.9 |
| PCR01 | 12 | 2 | 20 | Cyt b | 34.3 | 34.5 | 0.3 | 76.6 | 66.3 | 12.3 |
| PCR01 | 12 | 2 | 20 | Cyt b | 34.4 | 34.5 | 0.3 | 69.5 | 66.3 | 12.3 |
| PCR01 | 12 | 2 | 20 | Cyt b | 34.8 | 34.5 | 0.3 | 52.7 | 66.3 | 12.3 |
| PCR01 | 0 | FilterNC | | Cyt b | Undetermined | | | | | |
| PCR01 | 0 | FilterNC | | Cyt b | Undetermined | | | | | |
| PCR01 | 0 | FilterNC | | Cyt b | Undetermined | | | | | |
| PCR01 | PCRNC | | | Cyt b | Undetermined | | | | | |
| PCR01 | PCRNC | | | Cyt b | Undetermined | | | | | |
| PCR01 | PCRNC | | | Cyt b | Undetermined | | | | | |
| PCR02 | STANDARD | | | Cyt b | 25.4 | 25.4 | 0.1 | 30000.0 | 30000.0 | |
| PCR02 | STANDARD | | | Cyt b | 25.3 | 25.4 | 0.1 | 30000.0 | 30000.0 | |
| PCR02 | STANDARD | | | Cyt b | 25.4 | 25.4 | 0.1 | 30000.0 | 30000.0 | |
| PCR02 | STANDARD | | | Cyt b | 28.8 | 28.8 | 0.0 | 3000.0 | 3000.0 | |
| PCR02 | STANDARD | | | Cyt b | 28.8 | 28.8 | 0.0 | 3000.0 | 3000.0 | |
| PCR02 | STANDARD | | | Cyt b | 28.8 | 28.8 | 0.0 | 3000.0 | 3000.0 | |

| | | | | | | | | | | |
|---|---|---|---|---|---|---|---|---|---|---|
| PCR02 | STANDARD | | | Cyt b | 32.2 | 32.3 | 0.1 | 300.0 | 300.0 | |
| PCR02 | STANDARD | | | Cyt b | 32.4 | 32.3 | 0.1 | 300.0 | 300.0 | |
| PCR02 | STANDARD | | | Cyt b | 32.3 | 32.3 | 0.1 | 300.0 | 300.0 | |
| PCR02 | STANDARD | | | Cyt b | 35.9 | 35.8 | 0.2 | 30.0 | 30.0 | |
| PCR02 | STANDARD | | | Cyt b | 35.9 | 35.8 | 0.2 | 30.0 | 30.0 | |
| PCR02 | STANDARD | | | Cyt b | 35.6 | 35.8 | 0.2 | 30.0 | 30.0 | |
| PCR02 | 12 | 3 | 0.7 | Cyt b | 29.8 | 29.8 | 0.0 | 1547.8 | 1543.0 | 10.8 |
| PCR02 | 12 | 3 | 0.7 | Cyt b | 29.8 | 29.8 | 0.0 | 1550.6 | 1543.0 | 10.8 |
| PCR02 | 12 | 3 | 0.7 | Cyt b | 29.8 | 29.8 | 0.0 | 1530.6 | 1543.0 | 10.8 |
| PCR02 | 12 | 3 | 2 | Cyt b | 32.6 | 32.7 | 0.1 | 239.7 | 231.5 | 22.1 |
| PCR02 | 12 | 3 | 2 | Cyt b | 32.6 | 32.7 | 0.1 | 248.3 | 231.5 | 22.1 |
| PCR02 | 12 | 3 | 2 | Cyt b | 32.9 | 32.7 | 0.1 | 206.4 | 231.5 | 22.1 |
| PCR02 | 12 | 3 | 8 | Cyt b | Undetermined | 39.3 | 0.1 | | | |
| PCR02 | 12 | 3 | 8 | Cyt b | 39.4 | 39.3 | 0.1 | 2.7 | 2.9 | 0.2 |
| PCR02 | 12 | 3 | 8 | Cyt b | 39.2 | 39.3 | 0.1 | 3.0 | 2.9 | 0.2 |
| PCR02 | 12 | 3 | 14 | Cyt b | Undetermined | 38.0 | 0.4 | | | |
| PCR02 | 12 | 3 | 14 | Cyt b | 38.3 | 38.0 | 0.4 | 5.5 | 6.8 | 1.9 |
| PCR02 | 12 | 3 | 14 | Cyt b | 37.7 | 38.0 | 0.4 | 8.1 | 6.8 | 1.9 |
| PCR02 | 12 | 3 | 20 | Cyt b | 30.8 | 30.8 | 0.0 | 800.5 | 782.0 | 25.2 |
| PCR02 | 12 | 3 | 20 | Cyt b | 30.8 | 30.8 | 0.0 | 792.3 | 782.0 | 25.2 |
| PCR02 | 12 | 3 | 20 | Cyt b | 30.9 | 30.8 | 0.0 | 753.3 | 782.0 | 25.2 |
| PCR02 | 24 | 1 | 0.7 | Cyt b | 32.8 | 33.0 | 0.2 | 209.8 | 193.5 | 22.4 |
| PCR02 | 24 | 1 | 0.7 | Cyt b | 32.9 | 33.0 | 0.2 | 202.8 | 193.5 | 22.4 |
| PCR02 | 24 | 1 | 0.7 | Cyt b | 33.2 | 33.0 | 0.2 | 168.0 | 193.5 | 22.4 |
| PCR02 | 24 | 1 | 2 | Cyt b | Undetermined | | | | | |
| PCR02 | 24 | 1 | 2 | Cyt b | Undetermined | | | | | |
| PCR02 | 24 | 1 | 2 | Cyt b | Undetermined | | | | | |
| PCR02 | 24 | 1 | 8 | Cyt b | Undetermined | | | | | |
| PCR02 | 24 | 1 | 8 | Cyt b | Undetermined | | | | | |
| PCR02 | 24 | 1 | 8 | Cyt b | Undetermined | | | | | |
| PCR02 | 24 | 1 | 14 | Cyt b | Undetermined | | | | | |
| PCR02 | 24 | 1 | 14 | Cyt b | Undetermined | | | | | |
| PCR02 | 24 | 1 | 14 | Cyt b | Undetermined | | | | | |

| | | | | | | | | | | |
|---|---|---|---|---|---|---|---|---|---|---|
| PCR02 | 24 | 1 | 20 | Cyt b | 34.2 | 34.3 | 0.2 | 85.8 | 80.5 | 11.6 |
| PCR02 | 24 | 1 | 20 | Cyt b | 34.1 | 34.3 | 0.2 | 88.5 | 80.5 | 11.6 |
| PCR02 | 24 | 1 | 20 | Cyt b | 34.6 | 34.3 | 0.2 | 67.2 | 80.5 | 11.6 |
| PCR02 | 24 | 2 | 0.7 | Cyt b | 32.1 | 32.1 | 0.0 | 334.1 | 336.1 | 2.4 |
| PCR02 | 24 | 2 | 0.7 | Cyt b | 32.1 | 32.1 | 0.0 | 338.7 | 336.1 | 2.4 |
| PCR02 | 24 | 2 | 0.7 | Cyt b | 32.1 | 32.1 | 0.0 | 335.4 | 336.1 | 2.4 |
| PCR02 | 24 | 2 | 2 | Cyt b | Undetermined | | | | | |
| PCR02 | 24 | 2 | 2 | Cyt b | Undetermined | | | | | |
| PCR02 | 24 | 2 | 2 | Cyt b | Undetermined | | | | | |
| PCR02 | 24 | 2 | 8 | Cyt b | Undetermined | | | | | |
| PCR02 | 24 | 2 | 8 | Cyt b | Undetermined | | | | | |
| PCR02 | 24 | 2 | 8 | Cyt b | Undetermined | | | | | |
| PCR02 | 24 | 2 | 14 | Cyt b | Undetermined | | | | | |
| PCR02 | 24 | 2 | 14 | Cyt b | Undetermined | | | | | |
| PCR02 | 24 | 2 | 14 | Cyt b | Undetermined | | | | | |
| PCR02 | 24 | 2 | 20 | Cyt b | 39.4 | 38.7 | 1.0 | 2.7 | 4.8 | 3.0 |
| PCR02 | 24 | 2 | 20 | Cyt b | 38.0 | 38.7 | 1.0 | 7.0 | 4.8 | 3.0 |
| PCR02 | 24 | 2 | 20 | Cyt b | Undetermined | 38.7 | 1.0 | | | |
| PCR02 | 24 | 3 | 0.7 | Cyt b | 31.9 | 31.9 | 0.1 | 393.5 | 396.4 | 16.3 |
| PCR02 | 24 | 3 | 0.7 | Cyt b | 31.8 | 31.9 | 0.1 | 414.0 | 396.4 | 16.3 |
| PCR02 | 24 | 3 | 0.7 | Cyt b | 31.9 | 31.9 | 0.1 | 381.8 | 396.4 | 16.3 |
| PCR02 | 24 | 3 | 2 | Cyt b | Undetermined | | | | | |
| PCR02 | 24 | 3 | 2 | Cyt b | Undetermined | | | | | |
| PCR02 | 24 | 3 | 2 | Cyt b | Undetermined | | | | | |
| PCR02 | 24 | 3 | 8 | Cyt b | Undetermined | | | | | |
| PCR02 | 24 | 3 | 8 | Cyt b | Undetermined | | | | | |
| PCR02 | 24 | 3 | 8 | Cyt b | Undetermined | | | | | |
| PCR02 | 24 | 3 | 14 | Cyt b | Undetermined | | | | | |
| PCR02 | 24 | 3 | 14 | Cyt b | Undetermined | | | | | |
| PCR02 | 24 | 3 | 14 | Cyt b | Undetermined | | | | | |
| PCR02 | 24 | 3 | 20 | Cyt b | Undetermined | | | | | |
| PCR02 | 24 | 3 | 20 | Cyt b | Undetermined | | | | | |
| PCR02 | 24 | 3 | 20 | Cyt b | Undetermined | | | | | |

| | | | | | | | | | | |
|---|---|---|---|---|---|---|---|---|---|---|
| PCR02 | 12 | 1 | 20 | Cyt b | 40.0 | 39.6 | 0.3 | 1.9 | 2.4 | 0.5 |
| PCR02 | 12 | 1 | 20 | Cyt b | 39.4 | 39.6 | 0.3 | 2.7 | 2.4 | 0.5 |
| PCR02 | 12 | 1 | 20 | Cyt b | 39.4 | 39.6 | 0.3 | 2.6 | 2.4 | 0.5 |
| PCR02 | 12 | FilterNC | | Cyt b | Undetermined | | | | | |
| PCR02 | 12 | FilterNC | | Cyt b | Undetermined | | | | | |
| PCR02 | 12 | FilterNC | | Cyt b | Undetermined | | | | | |
| PCR02 | 24 | FilterNC | | Cyt b | Undetermined | | | | | |
| PCR02 | 24 | FilterNC | | Cyt b | Undetermined | | | | | |
| PCR02 | 24 | FilterNC | | Cyt b | Undetermined | | | | | |
| PCR02 | PCRNC | | | Cyt b | Undetermined | | | | | |
| PCR02 | PCRNC | | | Cyt b | Undetermined | | | | | |
| PCR02 | PCRNC | | | Cyt b | Undetermined | | | | | |
| PCR03 | STANDARD | | | ITS1 | 26.9 | 26.8 | 0.1 | 30000.0 | | |
| PCR03 | STANDARD | | | ITS1 | 26.8 | 26.8 | 0.1 | 30000.0 | | |
| PCR03 | STANDARD | | | ITS1 | 26.7 | 26.8 | 0.1 | 30000.0 | | |
| PCR03 | STANDARD | | | ITS1 | 30.5 | 30.5 | 0.0 | 3000.0 | | |
| PCR03 | STANDARD | | | ITS1 | 30.5 | 30.5 | 0.0 | 3000.0 | | |
| PCR03 | STANDARD | | | ITS1 | 30.5 | 30.5 | 0.0 | 3000.0 | | |
| PCR03 | STANDARD | | | ITS1 | 33.7 | 34.1 | 0.3 | 300.0 | | |
| PCR03 | STANDARD | | | ITS1 | 34.2 | 34.1 | 0.3 | 300.0 | | |
| PCR03 | STANDARD | | | ITS1 | 34.4 | 34.1 | 0.3 | 300.0 | | |
| PCR03 | STANDARD | | | ITS1 | 38.4 | 38.3 | 0.1 | 30.0 | | |
| PCR03 | STANDARD | | | ITS1 | 38.3 | 38.3 | 0.1 | 30.0 | | |
| PCR03 | STANDARD | | | ITS1 | 38.2 | 38.3 | 0.1 | 30.0 | | |
| PCR03 | 0 | 1 | 0.7 | ITS1 | 17.0 | 16.9 | 0.1 | 10548603.0 | 10935231.0 | 470061.6 |
| PCR03 | 0 | 1 | 0.7 | ITS1 | 16.8 | 16.9 | 0.1 | 11458464.0 | 10935231.0 | 470061.6 |
| PCR03 | 0 | 1 | 0.7 | ITS1 | 16.9 | 16.9 | 0.1 | 10798623.0 | 10935231.0 | 470061.6 |
| PCR03 | 0 | 1 | 2 | ITS1 | 27.8 | 27.8 | 0.0 | 15771.0 | 15411.0 | 362.2 |
| PCR03 | 0 | 1 | 2 | ITS1 | 27.8 | 27.8 | 0.0 | 15415.5 | 15411.0 | 362.2 |
| PCR03 | 0 | 1 | 2 | ITS1 | 27.8 | 27.8 | 0.0 | 15046.6 | 15411.0 | 362.2 |
| PCR03 | 0 | 1 | 8 | ITS1 | 29.8 | 29.8 | 0.1 | 4486.3 | 4735.7 | 285.5 |
| PCR03 | 0 | 1 | 8 | ITS1 | 29.7 | 29.8 | 0.1 | 5047.1 | 4735.7 | 285.5 |
| PCR03 | 0 | 1 | 8 | ITS1 | 29.8 | 29.8 | 0.1 | 4673.5 | 4735.7 | 285.5 |

| | | | | | | | | | | |
|---|---|---|---|---|---|---|---|---|---|---|
| PCR03 | 0 | 1 | 14 | ITS1 | 30.9 | 30.9 | 0.0 | 2329.5 | 2386.8 | 50.6 |
| PCR03 | 0 | 1 | 14 | ITS1 | 30.9 | 30.9 | 0.0 | 2405.8 | 2386.8 | 50.6 |
| PCR03 | 0 | 1 | 14 | ITS1 | 30.9 | 30.9 | 0.0 | 2425.3 | 2386.8 | 50.6 |
| PCR03 | 0 | 1 | 20 | ITS1 | 21.1 | 21.0 | 0.0 | 884475.8 | 913867.0 | 26159.2 |
| PCR03 | 0 | 1 | 20 | ITS1 | 21.0 | 21.0 | 0.0 | 922527.4 | 913867.0 | 26159.2 |
| PCR03 | 0 | 1 | 20 | ITS1 | 21.0 | 21.0 | 0.0 | 934597.8 | 913867.0 | 26159.2 |
| PCR03 | 0 | 2 | 0.7 | ITS1 | 17.7 | 17.7 | 0.1 | 6800620.5 | 6908624.5 | 258385.7 |
| PCR03 | 0 | 2 | 0.7 | ITS1 | 17.6 | 17.7 | 0.1 | 7203488.5 | 6908624.5 | 258385.7 |
| PCR03 | 0 | 2 | 0.7 | ITS1 | 17.7 | 17.7 | 0.1 | 6721764.5 | 6908624.5 | 258385.7 |
| PCR03 | 0 | 2 | 2 | ITS1 | 28.0 | 28.0 | 0.1 | 13455.7 | 13315.6 | 447.8 |
| PCR03 | 0 | 2 | 2 | ITS1 | 28.0 | 28.0 | 0.1 | 13676.7 | 13315.6 | 447.8 |
| PCR03 | 0 | 2 | 2 | ITS1 | 28.1 | 28.0 | 0.1 | 12814.6 | 13315.6 | 447.8 |
| PCR03 | 0 | 2 | 8 | ITS1 | 28.1 | 28.1 | 0.1 | 12600.8 | 12867.4 | 579.4 |
| PCR03 | 0 | 2 | 8 | ITS1 | 28.0 | 28.1 | 0.1 | 13532.1 | 12867.4 | 579.4 |
| PCR03 | 0 | 2 | 8 | ITS1 | 28.2 | 28.1 | 0.1 | 12469.2 | 12867.4 | 579.4 |
| PCR03 | 0 | 2 | 14 | ITS1 | 28.4 | 28.4 | 0.0 | 10656.0 | 10497.8 | 163.0 |
| PCR03 | 0 | 2 | 14 | ITS1 | 28.4 | 28.4 | 0.0 | 10507.0 | 10497.8 | 163.0 |
| PCR03 | 0 | 2 | 14 | ITS1 | 28.5 | 28.4 | 0.0 | 10330.4 | 10497.8 | 163.0 |
| PCR03 | 0 | 2 | 20 | ITS1 | 21.4 | 21.5 | 0.1 | 735808.2 | 709537.8 | 27066.3 |
| PCR03 | 0 | 2 | 20 | ITS1 | 21.4 | 21.5 | 0.1 | 711064.9 | 709537.8 | 27066.3 |
| PCR03 | 0 | 2 | 20 | ITS1 | 21.5 | 21.5 | 0.1 | 681740.3 | 709537.8 | 27066.3 |
| PCR03 | 0 | 3 | 0.7 | ITS1 | 18.3 | 18.3 | 0.0 | 4748273.0 | 4651131.0 | 90402.1 |
| PCR03 | 0 | 3 | 0.7 | ITS1 | 18.4 | 18.3 | 0.0 | 4569467.5 | 4651131.0 | 90402.1 |
| PCR03 | 0 | 3 | 0.7 | ITS1 | 18.3 | 18.3 | 0.0 | 4635652.0 | 4651131.0 | 90402.1 |
| PCR03 | 0 | 3 | 2 | ITS1 | 26.3 | 26.2 | 0.1 | 38841.6 | 39376.0 | 1263.2 |
| PCR03 | 0 | 3 | 2 | ITS1 | 26.2 | 26.2 | 0.1 | 40818.5 | 39376.0 | 1263.2 |
| PCR03 | 0 | 3 | 2 | ITS1 | 26.3 | 26.2 | 0.1 | 38467.8 | 39376.0 | 1263.2 |
| PCR03 | 0 | 3 | 8 | ITS1 | 31.2 | 31.1 | 0.1 | 1962.5 | 2065.5 | 98.1 |
| PCR03 | 0 | 3 | 8 | ITS1 | 31.1 | 31.1 | 0.1 | 2158.0 | 2065.5 | 98.1 |
| PCR03 | 0 | 3 | 8 | ITS1 | 31.1 | 31.1 | 0.1 | 2076.0 | 2065.5 | 98.1 |
| PCR03 | 0 | 3 | 14 | ITS1 | 28.3 | 28.3 | 0.0 | 11663.5 | 11474.0 | 334.8 |
| PCR03 | 0 | 3 | 14 | ITS1 | 28.3 | 28.3 | 0.0 | 11670.9 | 11474.0 | 334.8 |
| PCR03 | 0 | 3 | 14 | ITS1 | 28.3 | 28.3 | 0.0 | 11087.4 | 11474.0 | 334.8 |

| | | | | | | | | | | |
|---|---|---|---|---|---|---|---|---|---|---|
| PCR03 | 0 | 3 | 20 | ITS1 | 21.1 | 21.1 | 0.1 | 874075.1 | 901749.2 | 34377.6 |
| PCR03 | 0 | 3 | 20 | ITS1 | 21.1 | 21.1 | 0.1 | 890940.2 | 901749.2 | 34377.6 |
| PCR03 | 0 | 3 | 20 | ITS1 | 21.0 | 21.1 | 0.1 | 940232.2 | 901749.2 | 34377.6 |
| PCR03 | 0 | FilterNC | | ITS1 | Undetermined | | | | | |
| PCR03 | 0 | FilterNC | | ITS1 | Undetermined | | | | | |
| PCR03 | 0 | FilterNC | | ITS1 | Undetermined | | | | | |
| PCR03 | PCRNC | | | ITS1 | Undetermined | | | | | |
| PCR03 | PCRNC | | | ITS1 | Undetermined | | | | | |
| PCR03 | PCRNC | | | ITS1 | Undetermined | | | | | |
| PCR04 | STANDARD | | | ITS1 | 27.2 | 26.9 | 0.2 | 30000.0 | 30000.0 | |
| PCR04 | STANDARD | | | ITS1 | 26.9 | 26.9 | 0.2 | 30000.0 | 30000.0 | |
| PCR04 | STANDARD | | | ITS1 | 26.8 | 26.9 | 0.2 | 30000.0 | 30000.0 | |
| PCR04 | STANDARD | | | ITS1 | 30.9 | 30.9 | 0.1 | 3000.0 | 3000.0 | |
| PCR04 | STANDARD | | | ITS1 | 30.8 | 30.9 | 0.1 | 3000.0 | 3000.0 | |
| PCR04 | STANDARD | | | ITS1 | 30.8 | 30.9 | 0.1 | 3000.0 | 3000.0 | |
| PCR04 | STANDARD | | | ITS1 | 34.5 | 34.7 | 0.3 | 300.0 | 300.0 | |
| PCR04 | STANDARD | | | ITS1 | 34.5 | 34.7 | 0.3 | 300.0 | 300.0 | |
| PCR04 | STANDARD | | | ITS1 | 35.0 | 34.7 | 0.3 | 300.0 | 300.0 | |
| PCR04 | STANDARD | | | ITS1 | 38.8 | 38.4 | 0.5 | 30.0 | 30.0 | |
| PCR04 | STANDARD | | | ITS1 | 37.8 | 38.4 | 0.5 | 30.0 | 30.0 | |
| PCR04 | STANDARD | | | ITS1 | 38.6 | 38.4 | 0.5 | 30.0 | 30.0 | |
| PCR04 | 12 | 1 | 0.7 | ITS1 | 26.3 | 26.2 | 0.1 | 45424.9 | 48002.8 | 2465.7 |
| PCR04 | 12 | 1 | 0.7 | ITS1 | 26.1 | 26.2 | 0.1 | 50338.3 | 48002.8 | 2465.7 |
| PCR04 | 12 | 1 | 0.7 | ITS1 | 26.2 | 26.2 | 0.1 | 48245.3 | 48002.8 | 2465.7 |
| PCR04 | 12 | 1 | 2 | ITS1 | 31.9 | 31.8 | 0.1 | 1550.7 | 1649.0 | 116.1 |
| PCR04 | 12 | 1 | 2 | ITS1 | 31.8 | 31.8 | 0.1 | 1619.1 | 1649.0 | 116.1 |
| PCR04 | 12 | 1 | 2 | ITS1 | 31.7 | 31.8 | 0.1 | 1777.1 | 1649.0 | 116.1 |
| PCR04 | 12 | 1 | 8 | ITS1 | 33.1 | 33.0 | 0.1 | 733.1 | 786.4 | 46.2 |
| PCR04 | 12 | 1 | 8 | ITS1 | 33.0 | 33.0 | 0.1 | 814.9 | 786.4 | 46.2 |
| PCR04 | 12 | 1 | 8 | ITS1 | 33.0 | 33.0 | 0.1 | 811.3 | 786.4 | 46.2 |
| PCR04 | 12 | 1 | 14 | ITS1 | 38.6 | 40.4 | 1.8 | 27.7 | 13.3 | 12.9 |
| PCR04 | 12 | 1 | 14 | ITS1 | 42.3 | 40.4 | 1.8 | 3.0 | 13.3 | 12.9 |
| PCR04 | 12 | 1 | 14 | ITS1 | 40.4 | 40.4 | 1.8 | 9.1 | 13.3 | 12.9 |

| | | | | | | | | | | |
|---|---|---|---|---|---|---|---|---|---|---|
| PCR04 | 12 | 1 | 20 | ITS1 | 38.4 | 39.3 | 1.3 | 30.2 | 21.6 | 12.5 |
| PCR04 | 12 | 1 | 20 | ITS1 | 40.8 | 39.3 | 1.3 | 7.3 | 21.6 | 12.5 |
| PCR04 | 12 | 1 | 20 | ITS1 | 38.6 | 39.3 | 1.3 | 27.4 | 21.6 | 12.5 |
| PCR04 | 12 | 1 | 0.7 | ITS1 | 27.3 | 27.2 | 0.1 | 24122.7 | 25610.2 | 1288.3 |
| PCR04 | 12 | 1 | 0.7 | ITS1 | 27.2 | 27.2 | 0.1 | 26341.1 | 25610.2 | 1288.3 |
| PCR04 | 12 | 1 | 0.7 | ITS1 | 27.2 | 27.2 | 0.1 | 26366.7 | 25610.2 | 1288.3 |
| PCR04 | 12 | 1 | 2 | ITS1 | 34.3 | 34.2 | 0.2 | 369.9 | 384.5 | 51.6 |
| PCR04 | 12 | 1 | 2 | ITS1 | 34.0 | 34.2 | 0.2 | 441.8 | 384.5 | 51.6 |
| PCR04 | 12 | 1 | 2 | ITS1 | 34.4 | 34.2 | 0.2 | 341.8 | 384.5 | 51.6 |
| PCR04 | 12 | 1 | 8 | ITS1 | 34.3 | 34.1 | 0.2 | 372.2 | 419.2 | 45.2 |
| PCR04 | 12 | 1 | 8 | ITS1 | 34.1 | 34.1 | 0.2 | 422.8 | 419.2 | 45.2 |
| PCR04 | 12 | 1 | 8 | ITS1 | 33.9 | 34.1 | 0.2 | 462.5 | 419.2 | 45.2 |
| PCR04 | 12 | 1 | 14 | ITS1 | 35.0 | 35.3 | 0.3 | 244.3 | 206.5 | 34.5 |
| PCR04 | 12 | 1 | 14 | ITS1 | 35.5 | 35.3 | 0.3 | 176.9 | 206.5 | 34.5 |
| PCR04 | 12 | 1 | 14 | ITS1 | 35.3 | 35.3 | 0.3 | 198.2 | 206.5 | 34.5 |
| PCR04 | 12 | 1 | 20 | ITS1 | 34.4 | 34.6 | 0.3 | 354.8 | 307.1 | 57.1 |
| PCR04 | 12 | 1 | 20 | ITS1 | 34.5 | 34.6 | 0.3 | 322.7 | 307.1 | 57.1 |
| PCR04 | 12 | 1 | 20 | ITS1 | 35.0 | 34.6 | 0.3 | 243.8 | 307.1 | 57.1 |
| PCR04 | 12 | 1 | 0.7 | ITS1 | 27.3 | 27.3 | 0.0 | 24394.9 | 24377.8 | 67.3 |
| PCR04 | 12 | 1 | 0.7 | ITS1 | 27.3 | 27.3 | 0.0 | 24434.8 | 24377.8 | 67.3 |
| PCR04 | 12 | 1 | 0.7 | ITS1 | 27.3 | 27.3 | 0.0 | 24303.5 | 24377.8 | 67.3 |
| PCR04 | 12 | 1 | 2 | ITS1 | 31.9 | 32.0 | 0.1 | 1561.9 | 1444.6 | 123.5 |
| PCR04 | 12 | 1 | 2 | ITS1 | 32.0 | 32.0 | 0.1 | 1456.3 | 1444.6 | 123.5 |
| PCR04 | 12 | 1 | 2 | ITS1 | 32.2 | 32.0 | 0.1 | 1315.6 | 1444.6 | 123.5 |
| PCR04 | 12 | 1 | 8 | ITS1 | 34.9 | 35.6 | 0.6 | 247.8 | 175.9 | 63.8 |
| PCR04 | 12 | 1 | 8 | ITS1 | 36.1 | 35.6 | 0.6 | 126.2 | 175.9 | 63.8 |
| PCR04 | 12 | 1 | 8 | ITS1 | 35.7 | 35.6 | 0.6 | 153.7 | 175.9 | 63.8 |
| PCR04 | 12 | 1 | 14 | ITS1 | 37.1 | 37.7 | 0.6 | 66.4 | 48.1 | 16.8 |
| PCR04 | 12 | 1 | 14 | ITS1 | 38.3 | 37.7 | 0.6 | 33.3 | 48.1 | 16.8 |
| PCR04 | 12 | 1 | 14 | ITS1 | 37.8 | 37.7 | 0.6 | 44.6 | 48.1 | 16.8 |
| PCR04 | 12 | 1 | 20 | ITS1 | 30.7 | 30.8 | 0.1 | 3131.4 | 2936.1 | 171.8 |
| PCR04 | 12 | 1 | 20 | ITS1 | 30.9 | 30.8 | 0.1 | 2868.7 | 2936.1 | 171.8 |
| PCR04 | 12 | 1 | 20 | ITS1 | 30.9 | 30.8 | 0.1 | 2808.3 | 2936.1 | 171.8 |

| | | | | | | | | | | |
|---|---|---|---|---|---|---|---|---|---|---|
| PCR04 | 12 | FilterNC | | ITS1 | Undetermined | | | | | |
| PCR04 | 12 | FilterNC | | ITS1 | Undetermined | | | | | |
| PCR04 | 12 | FilterNC | | ITS1 | Undetermined | | | | | |
| PCR04 | PCRNC | | | ITS1 | Undetermined | | | | | |
| PCR04 | PCRNC | | | ITS1 | Undetermined | | | | | |
| PCR04 | PCRNC | | | ITS1 | Undetermined | | | | | |
| PCR05 | STANDARD | | | ITS1 | 27.3 | 27.1 | 0.1 | 30000.0 | 30000.0 | |
| PCR05 | STANDARD | | | ITS1 | 27.0 | 27.1 | 0.1 | 30000.0 | 30000.0 | |
| PCR05 | STANDARD | | | ITS1 | 27.1 | 27.1 | 0.1 | 30000.0 | 30000.0 | |
| PCR05 | STANDARD | | | ITS1 | 31.1 | 31.1 | 0.0 | 3000.0 | 3000.0 | |
| PCR05 | STANDARD | | | ITS1 | 31.1 | 31.1 | 0.0 | 3000.0 | 3000.0 | |
| PCR05 | STANDARD | | | ITS1 | 31.1 | 31.1 | 0.0 | 3000.0 | 3000.0 | |
| PCR05 | STANDARD | | | ITS1 | 34.9 | 34.9 | 0.3 | 300.0 | 300.0 | |
| PCR05 | STANDARD | | | ITS1 | 34.7 | 34.9 | 0.3 | 300.0 | 300.0 | |
| PCR05 | STANDARD | | | ITS1 | 35.2 | 34.9 | 0.3 | 300.0 | 300.0 | |
| PCR05 | STANDARD | | | ITS1 | 38.3 | 38.6 | 0.5 | 30.0 | 30.0 | |
| PCR05 | STANDARD | | | ITS1 | 38.5 | 38.6 | 0.5 | 30.0 | 30.0 | |
| PCR05 | STANDARD | | | ITS1 | 39.2 | 38.6 | 0.5 | 30.0 | 30.0 | |
| PCR05 | 24 | 1 | 0.7 | ITS1 | 30.7 | 30.5 | 0.1 | 3678.8 | 4045.2 | 318.7 |
| PCR05 | 24 | 1 | 0.7 | ITS1 | 30.4 | 30.5 | 0.1 | 4258.5 | 4045.2 | 318.7 |
| PCR05 | 24 | 1 | 0.7 | ITS1 | 30.5 | 30.5 | 0.1 | 4198.4 | 4045.2 | 318.7 |
| PCR05 | 24 | 1 | 2 | ITS1 | Undetermined | | | | | |
| PCR05 | 24 | 1 | 2 | ITS1 | Undetermined | | | | | |
| PCR05 | 24 | 1 | 2 | ITS1 | Undetermined | | | | | |
| PCR05 | 24 | 1 | 8 | ITS1 | Undetermined | | | | | |
| PCR05 | 24 | 1 | 8 | ITS1 | Undetermined | | | | | |
| PCR05 | 24 | 1 | 8 | ITS1 | Undetermined | | | | | |
| PCR05 | 24 | 1 | 14 | ITS1 | Undetermined | | | | | |
| PCR05 | 24 | 1 | 14 | ITS1 | Undetermined | | | | | |
| PCR05 | 24 | 1 | 14 | ITS1 | Undetermined | | | | | |
| PCR05 | 24 | 1 | 20 | ITS1 | 33.5 | 33.6 | 0.1 | 680.5 | 641.9 | 34.4 |
| PCR05 | 24 | 1 | 20 | ITS1 | 33.6 | 33.6 | 0.1 | 630.8 | 641.9 | 34.4 |
| PCR05 | 24 | 1 | 20 | ITS1 | 33.7 | 33.6 | 0.1 | 614.5 | 641.9 | 34.4 |

| | | | | | | | | | | |
|---|---|---|---|---|---|---|---|---|---|---|
| PCR05 | 24 | 2 | 0.7 | ITS1 | 30.1 | 30.1 | 0.1 | 5092.5 | 5342.2 | 255.2 |
| PCR05 | 24 | 2 | 0.7 | ITS1 | 30.1 | 30.1 | 0.1 | 5331.7 | 5342.2 | 255.2 |
| PCR05 | 24 | 2 | 0.7 | ITS1 | 30.0 | 30.1 | 0.1 | 5602.6 | 5342.2 | 255.2 |
| PCR05 | 24 | 2 | 2 | ITS1 | Undetermined | | | | | |
| PCR05 | 24 | 2 | 2 | ITS1 | Undetermined | | | | | |
| PCR05 | 24 | 2 | 2 | ITS1 | Undetermined | | | | | |
| PCR05 | 24 | 2 | 8 | ITS1 | Undetermined | | | | | |
| PCR05 | 24 | 2 | 8 | ITS1 | Undetermined | | | | | |
| PCR05 | 24 | 2 | 8 | ITS1 | Undetermined | | | | | |
| PCR05 | 24 | 2 | 14 | ITS1 | Undetermined | 43.5 | | | 0.0 | |
| PCR05 | 24 | 2 | 14 | ITS1 | 43.5 | 43.5 | | 1.7 | 1.7 | |
| PCR05 | 24 | 2 | 14 | ITS1 | Undetermined | 43.5 | | | 0.0 | |
| PCR05 | 24 | 2 | 20 | ITS1 | 38.1 | 40.4 | 2.0 | 41.8 | 17.6 | 21.0 |
| PCR05 | 24 | 2 | 20 | ITS1 | 42.0 | 40.4 | 2.0 | 4.2 | 17.6 | 21.0 |
| PCR05 | 24 | 2 | 20 | ITS1 | 41.1 | 40.4 | 2.0 | 7.0 | 17.6 | 21.0 |
| PCR05 | 24 | 3 | 0.7 | ITS1 | 28.5 | 28.5 | 0.0 | 13404.1 | 13158.2 | 217.2 |
| PCR05 | 24 | 3 | 0.7 | ITS1 | 28.6 | 28.5 | 0.0 | 13077.9 | 13158.2 | 217.2 |
| PCR05 | 24 | 3 | 0.7 | ITS1 | 28.6 | 28.5 | 0.0 | 12992.5 | 13158.2 | 217.2 |
| PCR05 | 24 | 3 | 2 | ITS1 | Undetermined | | | | | |
| PCR05 | 24 | 3 | 2 | ITS1 | Undetermined | | | | | |
| PCR05 | 24 | 3 | 2 | ITS1 | Undetermined | | | | | |
| PCR05 | 24 | 3 | 8 | ITS1 | Undetermined | | | | | |
| PCR05 | 24 | 3 | 8 | ITS1 | Undetermined | | | | | |
| PCR05 | 24 | 3 | 8 | ITS1 | Undetermined | | | | | |
| PCR05 | 24 | 3 | 14 | ITS1 | Undetermined | | | | | |
| PCR05 | 24 | 3 | 14 | ITS1 | Undetermined | | | | | |
| PCR05 | 24 | 3 | 14 | ITS1 | Undetermined | | | | | |
| PCR05 | 24 | 3 | 20 | ITS1 | Undetermined | | | | | |
| PCR05 | 24 | 3 | 20 | ITS1 | Undetermined | | | | | |
| PCR05 | 24 | 3 | 20 | ITS1 | Undetermined | | | | | |
| PCR05 | 24 | FilterNC | | ITS1 | Undetermined | | | | | |
| PCR05 | 24 | FilterNC | | ITS1 | Undetermined | | | | | |
| PCR05 | 24 | FilterNC | | ITS1 | Undetermined | | | | | |

| | | | |
|---|---|---|---|
| PCR05 | PCRNC | ITS1 | Undetermined |
| PCR05 | PCRNC | ITS1 | Undetermined |
| PCR05 | PCRNC | ITS1 | Undetermined |

Table S7 All raw data of qPCR (DNA copies per 2 μL template DNA) in Experiment 2.

| qPCR ID | Time point | Tank No. | Replication No. | Filter pore size | Region | Cᴛ | Cᴛ Mean | Cᴛ SD | Quantity | Quantity Mean | Quantity SD |
|---|---|---|---|---|---|---|---|---|---|---|---|
| PCR01 | STANDARD | | | | Cyt b | 25.3 | 25.2 | 0.1 | 30000.0 | 30000.0 | |
| PCR01 | STANDARD | | | | Cyt b | 25.2 | 25.2 | 0.1 | 30000.0 | 30000.0 | |
| PCR01 | STANDARD | | | | Cyt b | 25.2 | 25.2 | 0.1 | 30000.0 | 30000.0 | |
| PCR01 | STANDARD | | | | Cyt b | 28.6 | 28.5 | 0.1 | 3000.0 | 3000.0 | |
| PCR01 | STANDARD | | | | Cyt b | 28.5 | 28.5 | 0.1 | 3000.0 | 3000.0 | |
| PCR01 | STANDARD | | | | Cyt b | 28.5 | 28.5 | 0.1 | 3000.0 | 3000.0 | |
| PCR01 | STANDARD | | | | Cyt b | 31.9 | 31.8 | 0.1 | 300.0 | 300.0 | |
| PCR01 | STANDARD | | | | Cyt b | 31.7 | 31.8 | 0.1 | 300.0 | 300.0 | |
| PCR01 | STANDARD | | | | Cyt b | 31.9 | 31.8 | 0.1 | 300.0 | 300.0 | |
| PCR01 | STANDARD | | | | Cyt b | 34.9 | 35.0 | 0.2 | 30.0 | 30.0 | |
| PCR01 | STANDARD | | | | Cyt b | 35.2 | 35.0 | 0.2 | 30.0 | 30.0 | |
| PCR01 | STANDARD | | | | Cyt b | 35.0 | 35.0 | 0.2 | 30.0 | 30.0 | |
| PCR01 | sunrise | tank1 | 1 | 0.7 | Cyt b | 30.9 | 30.8 | 0.1 | 572.1 | 600.5 | 26.4 |
| PCR01 | sunrise | tank1 | 1 | 0.7 | Cyt b | 30.7 | 30.8 | 0.1 | 624.4 | 600.5 | 26.4 |
| PCR01 | sunrise | tank1 | 1 | 0.7 | Cyt b | 30.8 | 30.8 | 0.1 | 605.0 | 600.5 | 26.4 |
| PCR01 | sunrise | tank1 | 1 | 2 | Cyt b | 31.7 | 31.6 | 0.1 | 312.3 | 330.1 | 15.8 |
| PCR01 | sunrise | tank1 | 1 | 2 | Cyt b | 31.6 | 31.6 | 0.1 | 335.5 | 330.1 | 15.8 |
| PCR01 | sunrise | tank1 | 1 | 2 | Cyt b | 31.6 | 31.6 | 0.1 | 342.4 | 330.1 | 15.8 |
| PCR01 | sunrise | tank1 | 1 | 8 | Cyt b | 33.6 | 33.8 | 0.2 | 80.7 | 70.5 | 9.4 |
| PCR01 | sunrise | tank1 | 1 | 8 | Cyt b | 34.0 | 33.8 | 0.2 | 62.2 | 70.5 | 9.4 |
| PCR01 | sunrise | tank1 | 1 | 8 | Cyt b | 33.9 | 33.8 | 0.2 | 68.7 | 70.5 | 9.4 |
| PCR01 | sunrise | tank1 | 1 | 14 | Cyt b | 36.6 | 37.0 | 0.4 | 9.8 | 8.0 | 2.2 |
| PCR01 | sunrise | tank1 | 1 | 14 | Cyt b | 37.4 | 37.0 | 0.4 | 5.5 | 8.0 | 2.2 |
| PCR01 | sunrise | tank1 | 1 | 14 | Cyt b | 36.8 | 37.0 | 0.4 | 8.7 | 8.0 | 2.2 |
| PCR01 | sunrise | tank1 | 1 | 20 | Cyt b | 31.3 | 31.3 | 0.1 | 432.6 | 434.2 | 22.4 |
| PCR01 | sunrise | tank1 | 1 | 20 | Cyt b | 31.2 | 31.3 | 0.1 | 457.4 | 434.2 | 22.4 |
| PCR01 | sunrise | tank1 | 1 | 20 | Cyt b | 31.3 | 31.3 | 0.1 | 412.7 | 434.2 | 22.4 |
| PCR01 | sunrise | tank2 | 1 | 0.7 | Cyt b | 26.7 | 26.7 | 0.1 | 10598.8 | 11230.9 | 603.9 |
| PCR01 | sunrise | tank2 | 1 | 0.7 | Cyt b | 26.6 | 26.7 | 0.1 | 11292.2 | 11230.9 | 603.9 |
| PCR01 | sunrise | tank2 | 1 | 0.7 | Cyt b | 26.6 | 26.7 | 0.1 | 11801.8 | 11230.9 | 603.9 |
| PCR01 | sunrise | tank2 | 1 | 2 | Cyt b | 28.9 | 29.0 | 0.1 | 2313.1 | 2152.2 | 178.2 |
| PCR01 | sunrise | tank2 | 1 | 2 | Cyt b | 29.0 | 29.0 | 0.1 | 2182.9 | 2152.2 | 178.2 |
| PCR01 | sunrise | tank2 | 1 | 2 | Cyt b | 29.1 | 29.0 | 0.1 | 1960.6 | 2152.2 | 178.2 |

| | | | | | | | | | | | |
|---|---|---|---|---|---|---|---|---|---|---|---|
| PCR01 | sunrise | tank2 | 1 | 8 | Cyt b | 33.3 | 33.4 | 0.0 | 100.4 | 99.1 | 1.8 |
| PCR01 | sunrise | tank2 | 1 | 8 | Cyt b | 33.4 | 33.4 | 0.0 | 97.0 | 99.1 | 1.8 |
| PCR01 | sunrise | tank2 | 1 | 8 | Cyt b | 33.3 | 33.4 | 0.0 | 100.0 | 99.1 | 1.8 |
| PCR01 | sunrise | tank2 | 1 | 14 | Cyt b | 33.2 | 33.2 | 0.1 | 106.6 | 109.3 | 4.3 |
| PCR01 | sunrise | tank2 | 1 | 14 | Cyt b | 33.2 | 33.2 | 0.1 | 114.3 | 109.3 | 4.3 |
| PCR01 | sunrise | tank2 | 1 | 14 | Cyt b | 33.2 | 33.2 | 0.1 | 107.1 | 109.3 | 4.3 |
| PCR01 | sunrise | tank2 | 1 | 20 | Cyt b | 32.9 | 32.7 | 0.3 | 141.1 | 164.5 | 30.5 |
| PCR01 | sunrise | tank2 | 1 | 20 | Cyt b | 32.7 | 32.7 | 0.3 | 153.4 | 164.5 | 30.5 |
| PCR01 | sunrise | tank2 | 1 | 20 | Cyt b | 32.4 | 32.7 | 0.3 | 199.0 | 164.5 | 30.5 |
| PCR01 | PCRNC | | | | Cyt b | Undetermined | | | | | |
| PCR01 | PCRNC | | | | Cyt b | Undetermined | | | | | |
| PCR01 | PCRNC | | | | Cyt b | Undetermined | | | | | |
| PCR02 | STANDARD | | | | ITS1 | 26.9 | 26.8 | 0.1 | 30000.0 | | |
| PCR02 | STANDARD | | | | ITS1 | 26.8 | 26.8 | 0.1 | 30000.0 | | |
| PCR02 | STANDARD | | | | ITS1 | 26.7 | 26.8 | 0.1 | 30000.0 | | |
| PCR02 | STANDARD | | | | ITS1 | 30.5 | 30.5 | 0.0 | 3000.0 | | |
| PCR02 | STANDARD | | | | ITS1 | 30.5 | 30.5 | 0.0 | 3000.0 | | |
| PCR02 | STANDARD | | | | ITS1 | 30.5 | 30.5 | 0.0 | 3000.0 | | |
| PCR02 | STANDARD | | | | ITS1 | 33.7 | 34.1 | 0.3 | 300.0 | | |
| PCR02 | STANDARD | | | | ITS1 | 34.2 | 34.1 | 0.3 | 300.0 | | |
| PCR02 | STANDARD | | | | ITS1 | 34.4 | 34.1 | 0.3 | 300.0 | | |
| PCR02 | STANDARD | | | | ITS1 | 38.4 | 38.3 | 0.1 | 30.0 | | |
| PCR02 | STANDARD | | | | ITS1 | 38.3 | 38.3 | 0.1 | 30.0 | | |
| PCR02 | STANDARD | | | | ITS1 | 38.2 | 38.3 | 0.1 | 30.0 | | |
| PCR02 | sunrise | tank1 | 1 | 0.7 | ITS1 | 29.4 | 29.4 | 0.1 | 5816.1 | 6016.9 | 195.1 |
| PCR02 | sunrise | tank1 | 1 | 0.7 | ITS1 | 29.3 | 29.4 | 0.1 | 6205.8 | 6016.9 | 195.1 |
| PCR02 | sunrise | tank1 | 1 | 0.7 | ITS1 | 29.4 | 29.4 | 0.1 | 6028.9 | 6016.9 | 195.1 |
| PCR02 | sunrise | tank1 | 1 | 2 | ITS1 | 33.4 | 33.3 | 0.2 | 529.1 | 575.3 | 81.3 |
| PCR02 | sunrise | tank1 | 1 | 2 | ITS1 | 33.0 | 33.3 | 0.2 | 669.1 | 575.3 | 81.3 |
| PCR02 | sunrise | tank1 | 1 | 2 | ITS1 | 33.4 | 33.3 | 0.2 | 527.6 | 575.3 | 81.3 |
| PCR02 | sunrise | tank1 | 1 | 8 | ITS1 | 36.0 | 35.4 | 0.5 | 110.8 | 158.1 | 47.0 |
| PCR02 | sunrise | tank1 | 1 | 8 | ITS1 | 35.0 | 35.4 | 0.5 | 204.6 | 158.1 | 47.0 |
| PCR02 | sunrise | tank1 | 1 | 8 | ITS1 | 35.4 | 35.4 | 0.5 | 159.0 | 158.1 | 47.0 |
| PCR02 | sunrise | tank1 | 1 | 14 | ITS1 | 37.9 | 37.6 | 0.7 | 35.4 | 44.7 | 20.4 |
| PCR02 | sunrise | tank1 | 1 | 14 | ITS1 | 38.1 | 37.6 | 0.7 | 30.7 | 44.7 | 20.4 |
| PCR02 | sunrise | tank1 | 1 | 14 | ITS1 | 36.8 | 37.6 | 0.7 | 68.1 | 44.7 | 20.4 |

| | | | | | | | | | | | |
|---|---|---|---|---|---|---|---|---|---|---|---|
| PCR02 | sunrise | tank1 | 1 | 20 | ITS1 | 34.6 | 34.6 | 0.1 | 248.7 | 258.1 | 13.5 |
| PCR02 | sunrise | tank1 | 1 | 20 | ITS1 | 34.6 | 34.6 | 0.1 | 252.0 | 258.1 | 13.5 |
| PCR02 | sunrise | tank1 | 1 | 20 | ITS1 | 34.5 | 34.6 | 0.1 | 273.5 | 258.1 | 13.5 |
| PCR02 | sunrise | tank2 | 1 | 0.7 | ITS1 | 24.7 | 24.7 | 0.0 | 101163.3 | 103073.0 | 1662.3 |
| PCR02 | sunrise | tank2 | 1 | 0.7 | ITS1 | 24.6 | 24.7 | 0.0 | 104194.9 | 103073.0 | 1662.3 |
| PCR02 | sunrise | tank2 | 1 | 0.7 | ITS1 | 24.6 | 24.7 | 0.0 | 103860.8 | 103073.0 | 1662.3 |
| PCR02 | sunrise | tank2 | 1 | 2 | ITS1 | 30.0 | 30.0 | 0.0 | 4137.2 | 4130.5 | 6.2 |
| PCR02 | sunrise | tank2 | 1 | 2 | ITS1 | 30.0 | 30.0 | 0.0 | 4125.0 | 4130.5 | 6.2 |
| PCR02 | sunrise | tank2 | 1 | 2 | ITS1 | 30.0 | 30.0 | 0.0 | 4129.2 | 4130.5 | 6.2 |
| PCR02 | sunrise | tank2 | 1 | 8 | ITS1 | 32.1 | 32.0 | 0.1 | 1119.4 | 1212.1 | 88.1 |
| PCR02 | sunrise | tank2 | 1 | 8 | ITS1 | 31.9 | 32.0 | 0.1 | 1294.8 | 1212.1 | 88.1 |
| PCR02 | sunrise | tank2 | 1 | 8 | ITS1 | 32.0 | 32.0 | 0.1 | 1222.0 | 1212.1 | 88.1 |
| PCR02 | sunrise | tank2 | 1 | 20 | ITS1 | 33.8 | 33.7 | 0.1 | 424.0 | 450.7 | 33.4 |
| PCR02 | sunrise | tank2 | 1 | 20 | ITS1 | 33.5 | 33.7 | 0.1 | 488.1 | 450.7 | 33.4 |
| PCR02 | sunrise | tank2 | 1 | 20 | ITS1 | 33.7 | 33.7 | 0.1 | 440.1 | 450.7 | 33.4 |
| PCR02 | sunrise | tank2 | 1 | 14 | ITS1 | 33.0 | 33.1 | 0.1 | 666.8 | 641.0 | 26.2 |
| PCR02 | sunrise | tank2 | 1 | 14 | ITS1 | 33.1 | 33.1 | 0.1 | 641.8 | 641.0 | 26.2 |
| PCR02 | sunrise | tank2 | 1 | 14 | ITS1 | 33.1 | 33.1 | 0.1 | 614.4 | 641.0 | 26.2 |
| PCR02 | PCRNC | | | | ITS1 | Undetermined | | | | | |
| PCR02 | PCRNC | | | | ITS1 | Undetermined | | | | | |
| PCR02 | PCRNC | | | | ITS1 | Undetermined | | | | | |
| PCR03 | STANDARD | | | | Cyt b | 25.6 | 25.4 | 0.1 | 30000.0 | | |
| PCR03 | STANDARD | | | | Cyt b | 25.4 | 25.4 | 0.1 | 30000.0 | | |
| PCR03 | STANDARD | | | | Cyt b | 25.4 | 25.4 | 0.1 | 30000.0 | | |
| PCR03 | STANDARD | | | | Cyt b | 28.9 | 28.9 | 0.1 | 3000.0 | | |
| PCR03 | STANDARD | | | | Cyt b | 28.8 | 28.9 | 0.1 | 3000.0 | | |
| PCR03 | STANDARD | | | | Cyt b | 28.9 | 28.9 | 0.1 | 3000.0 | | |
| PCR03 | STANDARD | | | | Cyt b | 32.4 | 32.4 | 0.1 | 300.0 | | |
| PCR03 | STANDARD | | | | Cyt b | 32.4 | 32.4 | 0.1 | 300.0 | | |
| PCR03 | STANDARD | | | | Cyt b | 32.3 | 32.4 | 0.1 | 300.0 | | |
| PCR03 | STANDARD | | | | Cyt b | 35.4 | 35.4 | 0.2 | 30.0 | | |
| PCR03 | STANDARD | | | | Cyt b | 35.6 | 35.4 | 0.2 | 30.0 | | |
| PCR03 | STANDARD | | | | Cyt b | 35.2 | 35.4 | 0.2 | 30.0 | | |
| PCR03 | sunrise | tank2 | 2 | 0.7 | Cyt b | 26.7 | 26.7 | 0.1 | 12892.8 | 13681.3 | 918.6 |
| PCR03 | sunrise | tank2 | 2 | 0.7 | Cyt b | 26.7 | 26.7 | 0.1 | 13461.1 | 13681.3 | 918.6 |
| PCR03 | sunrise | tank2 | 2 | 0.7 | Cyt b | 26.5 | 26.7 | 0.1 | 14689.9 | 13681.3 | 918.6 |

| | | | | | | | | | | | |
|---|---|---|---|---|---|---|---|---|---|---|---|
| PCR03 | sunrise | tank2 | 2 | 2 | Cyt b | 29.4 | 29.5 | 0.1 | 2096.9 | 1899.3 | 173.0 |
| PCR03 | sunrise | tank2 | 2 | 2 | Cyt b | 29.6 | 29.5 | 0.1 | 1826.4 | 1899.3 | 173.0 |
| PCR03 | sunrise | tank2 | 2 | 2 | Cyt b | 29.6 | 29.5 | 0.1 | 1774.8 | 1899.3 | 173.0 |
| PCR03 | sunrise | tank2 | 2 | 8 | Cyt b | 33.7 | 33.4 | 0.3 | 105.8 | 128.9 | 29.4 |
| PCR03 | sunrise | tank2 | 2 | 8 | Cyt b | 33.1 | 33.4 | 0.3 | 162.0 | 128.9 | 29.4 |
| PCR03 | sunrise | tank2 | 2 | 8 | Cyt b | 33.5 | 33.4 | 0.3 | 119.0 | 128.9 | 29.4 |
| PCR03 | sunrise | tank2 | 2 | 14 | Cyt b | 32.8 | 32.8 | 0.2 | 196.8 | 203.1 | 30.9 |
| PCR03 | sunrise | tank2 | 2 | 14 | Cyt b | 33.0 | 32.8 | 0.2 | 175.8 | 203.1 | 30.9 |
| PCR03 | sunrise | tank2 | 2 | 14 | Cyt b | 32.5 | 32.8 | 0.2 | 236.6 | 203.1 | 30.9 |
| PCR03 | sunrise | tank2 | 2 | 20 | Cyt b | 30.2 | 30.2 | 0.1 | 1166.8 | 1217.9 | 65.5 |
| PCR03 | sunrise | tank2 | 2 | 20 | Cyt b | 30.2 | 30.2 | 0.1 | 1195.1 | 1217.9 | 65.5 |
| PCR03 | sunrise | tank2 | 2 | 20 | Cyt b | 30.1 | 30.2 | 0.1 | 1291.8 | 1217.9 | 65.5 |
| PCR03 | PCRNC | | | | Cyt b | Undetermined | | | | | |
| PCR03 | PCRNC | | | | Cyt b | Undetermined | | | | | |
| PCR03 | PCRNC | | | | Cyt b | Undetermined | | | | | |
| PCR04 | STANDARD | | | | ITS1 | 27.4 | 27.4 | 0.1 | 30000.0 | | |
| PCR04 | STANDARD | | | | ITS1 | 27.3 | 27.4 | 0.1 | 30000.0 | | |
| PCR04 | STANDARD | | | | ITS1 | 27.4 | 27.4 | 0.1 | 30000.0 | | |
| PCR04 | STANDARD | | | | ITS1 | 31.3 | 31.2 | 0.1 | 3000.0 | | |
| PCR04 | STANDARD | | | | ITS1 | 31.1 | 31.2 | 0.1 | 3000.0 | | |
| PCR04 | STANDARD | | | | ITS1 | 31.1 | 31.2 | 0.1 | 3000.0 | | |
| PCR04 | STANDARD | | | | ITS1 | 35.1 | 35.2 | 0.3 | 300.0 | | |
| PCR04 | STANDARD | | | | ITS1 | 35.0 | 35.2 | 0.3 | 300.0 | | |
| PCR04 | STANDARD | | | | ITS1 | 35.5 | 35.2 | 0.3 | 300.0 | | |
| PCR04 | STANDARD | | | | ITS1 | 39.0 | 38.9 | 0.1 | 30.0 | | |
| PCR04 | STANDARD | | | | ITS1 | 38.8 | 38.9 | 0.1 | 30.0 | | |
| PCR04 | STANDARD | | | | ITS1 | 38.9 | 38.9 | 0.1 | 30.0 | | |
| PCR04 | sunrise | tank2 | 2 | 0.7 | ITS1 | 24.9 | 24.9 | 0.0 | 133462.5 | 133406.6 | 1673.6 |
| PCR04 | sunrise | tank2 | 2 | 0.7 | ITS1 | 24.8 | 24.9 | 0.0 | 135051.6 | 133406.6 | 1673.6 |
| PCR04 | sunrise | tank2 | 2 | 0.7 | ITS1 | 24.9 | 24.9 | 0.0 | 131705.8 | 133406.6 | 1673.6 |
| PCR04 | sunrise | tank2 | 2 | 2 | ITS1 | 31.0 | 30.9 | 0.1 | 3381.4 | 3558.3 | 158.2 |
| PCR04 | sunrise | tank2 | 2 | 2 | ITS1 | 30.9 | 30.9 | 0.1 | 3607.2 | 3558.3 | 158.2 |
| PCR04 | sunrise | tank2 | 2 | 2 | ITS1 | 30.9 | 30.9 | 0.1 | 3686.3 | 3558.3 | 158.2 |
| PCR04 | sunrise | tank2 | 2 | 8 | ITS1 | 33.6 | 33.7 | 0.2 | 745.1 | 706.8 | 72.6 |
| PCR04 | sunrise | tank2 | 2 | 8 | ITS1 | 33.5 | 33.7 | 0.2 | 752.1 | 706.8 | 72.6 |
| PCR04 | sunrise | tank2 | 2 | 8 | ITS1 | 33.9 | 33.7 | 0.2 | 623.1 | 706.8 | 72.6 |

| | | | | | | | | | | | |
|---|---|---|---|---|---|---|---|---|---|---|---|
| PCR04 | sunrise | tank2 | 2 | 14 | ITS1 | 34.0 | 33.6 | 0.4 | 583.0 | 759.2 | 170.4 |
| PCR04 | sunrise | tank2 | 2 | 14 | ITS1 | 33.2 | 33.6 | 0.4 | 923.1 | 759.2 | 170.4 |
| PCR04 | sunrise | tank2 | 2 | 14 | ITS1 | 33.5 | 33.6 | 0.4 | 771.5 | 759.2 | 170.4 |
| PCR04 | sunrise | tank2 | 2 | 20 | ITS1 | 31.7 | 31.6 | 0.2 | 2217.8 | 2475.6 | 224.2 |
| PCR04 | sunrise | tank2 | 2 | 20 | ITS1 | 31.5 | 31.6 | 0.2 | 2624.8 | 2475.6 | 224.2 |
| PCR04 | sunrise | tank2 | 2 | 20 | ITS1 | 31.5 | 31.6 | 0.2 | 2584.3 | 2475.6 | 224.2 |
| PCR04 | PCRNC | | | | ITS1 | Undetermined | | | | | |
| PCR04 | PCRNC | | | | ITS1 | Undetermined | | | | | |
| PCR04 | PCRNC | | | | ITS1 | Undetermined | | | | | |
| PCR05 | STANDARD | | | | Cyt b | 25.3 | 25.2 | 0.1 | 30000.0 | | |
| PCR05 | STANDARD | | | | Cyt b | 25.2 | 25.2 | 0.1 | 30000.0 | | |
| PCR05 | STANDARD | | | | Cyt b | 25.1 | 25.2 | 0.1 | 30000.0 | | |
| PCR05 | STANDARD | | | | Cyt b | 28.9 | 29.0 | 0.0 | 3000.0 | | |
| PCR05 | STANDARD | | | | Cyt b | 29.0 | 29.0 | 0.0 | 3000.0 | | |
| PCR05 | STANDARD | | | | Cyt b | 29.0 | 29.0 | 0.0 | 3000.0 | | |
| PCR05 | STANDARD | | | | Cyt b | 31.9 | 32.1 | 0.1 | 300.0 | | |
| PCR05 | STANDARD | | | | Cyt b | 32.2 | 32.1 | 0.1 | 300.0 | | |
| PCR05 | STANDARD | | | | Cyt b | 32.2 | 32.1 | 0.1 | 300.0 | | |
| PCR05 | STANDARD | | | | Cyt b | 36.1 | 36.2 | 0.2 | 30.0 | | |
| PCR05 | STANDARD | | | | Cyt b | 36.1 | 36.2 | 0.2 | 30.0 | | |
| PCR05 | STANDARD | | | | Cyt b | 36.5 | 36.2 | 0.2 | 30.0 | | |
| PCR05 | sunrise | tank1 | 2 | 0.7 | Cyt b | 30.1 | 30.0 | 0.1 | 1306.9 | 1429.4 | 107.1 |
| PCR05 | sunrise | tank1 | 2 | 0.7 | Cyt b | 29.9 | 30.0 | 0.1 | 1476.5 | 1429.4 | 107.1 |
| PCR05 | sunrise | tank1 | 2 | 0.7 | Cyt b | 29.9 | 30.0 | 0.1 | 1504.9 | 1429.4 | 107.1 |
| PCR05 | sunrise | tank1 | 2 | 2 | Cyt b | 32.3 | 32.2 | 0.1 | 326.8 | 357.2 | 31.3 |
| PCR05 | sunrise | tank1 | 2 | 2 | Cyt b | 32.0 | 32.2 | 0.1 | 389.3 | 357.2 | 31.3 |
| PCR05 | sunrise | tank1 | 2 | 2 | Cyt b | 32.2 | 32.2 | 0.1 | 355.4 | 357.2 | 31.3 |
| PCR05 | sunrise | tank1 | 2 | 8 | Cyt b | 35.0 | 35.1 | 0.3 | 59.5 | 56.8 | 11.2 |
| PCR05 | sunrise | tank1 | 2 | 8 | Cyt b | 35.4 | 35.1 | 0.3 | 44.5 | 56.8 | 11.2 |
| PCR05 | sunrise | tank1 | 2 | 8 | Cyt b | 34.8 | 35.1 | 0.3 | 66.2 | 56.8 | 11.2 |
| PCR05 | sunrise | tank1 | 2 | 14 | Cyt b | 35.7 | 37.2 | 1.5 | 37.0 | 19.0 | 16.2 |
| PCR05 | sunrise | tank1 | 2 | 14 | Cyt b | 37.2 | 37.2 | 1.5 | 14.9 | 19.0 | 16.2 |
| PCR05 | sunrise | tank1 | 2 | 14 | Cyt b | 38.8 | 37.2 | 1.5 | 5.3 | 19.0 | 16.2 |
| PCR05 | sunrise | tank1 | 2 | 20 | Cyt b | 31.3 | 31.3 | 0.0 | 623.6 | 625.1 | 2.8 |
| PCR05 | sunrise | tank1 | 2 | 20 | Cyt b | 31.3 | 31.3 | 0.0 | 628.3 | 625.1 | 2.8 |
| PCR05 | sunrise | tank1 | 2 | 20 | Cyt b | 31.3 | 31.3 | 0.0 | 623.5 | 625.1 | 2.8 |

| | | | | | | | | | | | |
|---|---|---|---|---|---|---|---|---|---|---|---|
| PCR05 | sunrise | tank1 | 3 | 0.7 | Cyt b | 30.0 | 29.9 | 0.1 | 1431.8 | 1468.7 | 107.5 |
| PCR05 | sunrise | tank1 | 3 | 0.7 | Cyt b | 30.0 | 29.9 | 0.1 | 1384.6 | 1468.7 | 107.5 |
| PCR05 | sunrise | tank1 | 3 | 0.7 | Cyt b | 29.8 | 29.9 | 0.1 | 1589.8 | 1468.7 | 107.5 |
| PCR05 | sunrise | tank1 | 3 | 2 | Cyt b | 32.0 | 32.1 | 0.1 | 400.2 | 373.8 | 33.5 |
| PCR05 | sunrise | tank1 | 3 | 2 | Cyt b | 32.3 | 32.1 | 0.1 | 336.1 | 373.8 | 33.5 |
| PCR05 | sunrise | tank1 | 3 | 2 | Cyt b | 32.0 | 32.1 | 0.1 | 385.2 | 373.8 | 33.5 |
| PCR05 | sunrise | tank1 | 3 | 8 | Cyt b | 35.2 | 35.2 | 0.1 | 52.6 | 51.4 | 4.7 |
| PCR05 | sunrise | tank1 | 3 | 8 | Cyt b | 35.4 | 35.2 | 0.1 | 46.3 | 51.4 | 4.7 |
| PCR05 | sunrise | tank1 | 3 | 8 | Cyt b | 35.1 | 35.2 | 0.1 | 55.4 | 51.4 | 4.7 |
| PCR05 | sunrise | tank1 | 3 | 14 | Cyt b | 37.6 | 37.3 | 0.2 | 11.5 | 13.8 | 2.0 |
| PCR05 | sunrise | tank1 | 3 | 14 | Cyt b | 37.2 | 37.3 | 0.2 | 14.8 | 13.8 | 2.0 |
| PCR05 | sunrise | tank1 | 3 | 14 | Cyt b | 37.1 | 37.3 | 0.2 | 15.1 | 13.8 | 2.0 |
| PCR05 | sunrise | tank1 | 3 | 20 | Cyt b | 31.6 | 31.5 | 0.2 | 504.7 | 538.5 | 81.7 |
| PCR05 | sunrise | tank1 | 3 | 20 | Cyt b | 31.7 | 31.5 | 0.2 | 479.1 | 538.5 | 81.7 |
| PCR05 | sunrise | tank1 | 3 | 20 | Cyt b | 31.3 | 31.5 | 0.2 | 631.6 | 538.5 | 81.7 |
| PCR05 | sunset | tank1 | 1 | 0.7 | Cyt b | 30.9 | 30.9 | 0.0 | 811.2 | 813.1 | 16.9 |
| PCR05 | sunset | tank1 | 1 | 0.7 | Cyt b | 30.8 | 30.9 | 0.0 | 830.9 | 813.1 | 16.9 |
| PCR05 | sunset | tank1 | 1 | 0.7 | Cyt b | 30.9 | 30.9 | 0.0 | 797.1 | 813.1 | 16.9 |
| PCR05 | sunset | tank1 | 1 | 2 | Cyt b | 32.0 | 32.0 | 0.0 | 400.7 | 405.4 | 9.5 |
| PCR05 | sunset | tank1 | 1 | 2 | Cyt b | 31.9 | 32.0 | 0.0 | 416.3 | 405.4 | 9.5 |
| PCR05 | sunset | tank1 | 1 | 2 | Cyt b | 32.0 | 32.0 | 0.0 | 399.1 | 405.4 | 9.5 |
| PCR05 | sunset | tank1 | 1 | 8 | Cyt b | 35.0 | 34.9 | 0.2 | 58.1 | 63.2 | 9.3 |
| PCR05 | sunset | tank1 | 1 | 8 | Cyt b | 35.0 | 34.9 | 0.2 | 57.4 | 63.2 | 9.3 |
| PCR05 | sunset | tank1 | 1 | 8 | Cyt b | 34.6 | 34.9 | 0.2 | 74.0 | 63.2 | 9.3 |
| PCR05 | sunset | tank1 | 1 | 14 | Cyt b | 35.9 | 35.5 | 0.3 | 33.2 | 42.7 | 8.2 |
| PCR05 | sunset | tank1 | 1 | 14 | Cyt b | 35.3 | 35.5 | 0.3 | 46.9 | 42.7 | 8.2 |
| PCR05 | sunset | tank1 | 1 | 14 | Cyt b | 35.3 | 35.5 | 0.3 | 47.9 | 42.7 | 8.2 |
| PCR05 | sunset | tank1 | 1 | 20 | Cyt b | 34.6 | 34.6 | 0.0 | 77.4 | 76.6 | 0.8 |
| PCR05 | sunset | tank1 | 1 | 20 | Cyt b | 34.6 | 34.6 | 0.0 | 76.8 | 76.6 | 0.8 |
| PCR05 | sunset | tank1 | 1 | 20 | Cyt b | 34.6 | 34.6 | 0.0 | 75.8 | 76.6 | 0.8 |
| PCR05 | sunrise | tank2 | 3 | 0.7 | Cyt b | 27.0 | 27.0 | 0.0 | 9350.2 | 9519.5 | 218.4 |
| PCR05 | sunrise | tank2 | 3 | 0.7 | Cyt b | 27.0 | 27.0 | 0.0 | 9766.0 | 9519.5 | 218.4 |
| PCR05 | sunrise | tank2 | 3 | 0.7 | Cyt b | 27.0 | 27.0 | 0.0 | 9442.2 | 9519.5 | 218.4 |
| PCR05 | sunrise | tank2 | 3 | 2 | Cyt b | 29.6 | 29.6 | 0.1 | 1837.0 | 1858.8 | 79.4 |
| PCR05 | sunrise | tank2 | 3 | 2 | Cyt b | 29.6 | 29.6 | 0.1 | 1792.5 | 1858.8 | 79.4 |
| PCR05 | sunrise | tank2 | 3 | 2 | Cyt b | 29.5 | 29.6 | 0.1 | 1946.8 | 1858.8 | 79.4 |

| | | | | | | | | | | | |
|---|---|---|---|---|---|---|---|---|---|---|---|
| PCR05 | sunrise | tank2 | 3 | 8 | Cyt b | 34.6 | 34.5 | 0.1 | 74.1 | 79.0 | 5.4 |
| PCR05 | sunrise | tank2 | 3 | 8 | Cyt b | 34.4 | 34.5 | 0.1 | 84.8 | 79.0 | 5.4 |
| PCR05 | sunrise | tank2 | 3 | 8 | Cyt b | 34.5 | 34.5 | 0.1 | 78.1 | 79.0 | 5.4 |
| PCR05 | sunrise | tank2 | 3 | 14 | Cyt b | 34.0 | 34.2 | 0.3 | 114.1 | 97.0 | 17.7 |
| PCR05 | sunrise | tank2 | 3 | 14 | Cyt b | 34.5 | 34.2 | 0.3 | 78.8 | 97.0 | 17.7 |
| PCR05 | sunrise | tank2 | 3 | 14 | Cyt b | 34.2 | 34.2 | 0.3 | 98.2 | 97.0 | 17.7 |
| PCR05 | sunrise | tank2 | 3 | 20 | Cyt b | 30.6 | 30.6 | 0.0 | 960.3 | 990.4 | 26.3 |
| PCR05 | sunrise | tank2 | 3 | 20 | Cyt b | 30.5 | 30.6 | 0.0 | 1001.6 | 990.4 | 26.3 |
| PCR05 | sunrise | tank2 | 3 | 20 | Cyt b | 30.5 | 30.6 | 0.0 | 1009.3 | 990.4 | 26.3 |
| PCR05 | sunset | inlet seawater | | | Cyt b | Undetermined | | | | | |
| PCR05 | sunset | inlet seawater | | | Cyt b | Undetermined | | | | | |
| PCR05 | sunset | inlet seawater | | | Cyt b | Undetermined | | | | | |
| PCR05 | sunrise | inlet seawater | | | Cyt b | Undetermined | 39.6 | 1.0 | 0.0 | 2.3 | 2.5 |
| PCR05 | sunrise | inlet seawater | | | Cyt b | 38.9 | 39.6 | 1.0 | 5.1 | 2.3 | 2.5 |
| PCR05 | sunrise | inlet seawater | | | Cyt b | 40.3 | 39.6 | 1.0 | 2.0 | 2.3 | 2.5 |
| PCR05 | sunset | FilterNC | | | Cyt b | Undetermined | | | | | |
| PCR05 | sunset | FilterNC | | | Cyt b | Undetermined | | | | | |
| PCR05 | sunset | FilterNC | | | Cyt b | Undetermined | | | | | |
| PCR05 | sunrise | FilterNC | | | Cyt b | Undetermined | | | | | |
| PCR05 | sunrise | FilterNC | | | Cyt b | Undetermined | | | | | |
| PCR05 | sunrise | FilterNC | | | Cyt b | Undetermined | | | | | |
| PCR05 | PCRNC | | | | Cyt b | Undetermined | | | | | |
| PCR05 | PCRNC | | | | Cyt b | Undetermined | | | | | |
| PCR05 | PCRNC | | | | Cyt b | Undetermined | | | | | |
| PCR06 | STANDARD | | | | ITS1 | 27.3 | 27.1 | 0.2 | 30000.0 | | |
| PCR06 | STANDARD | | | | ITS1 | 27.0 | 27.1 | 0.2 | 30000.0 | | |
| PCR06 | STANDARD | | | | ITS1 | 27.0 | 27.1 | 0.2 | 30000.0 | | |
| PCR06 | STANDARD | | | | ITS1 | 31.3 | 31.2 | 0.1 | 3000.0 | | |
| PCR06 | STANDARD | | | | ITS1 | 31.2 | 31.2 | 0.1 | 3000.0 | | |
| PCR06 | STANDARD | | | | ITS1 | 31.2 | 31.2 | 0.1 | 3000.0 | | |
| PCR06 | STANDARD | | | | ITS1 | 34.7 | 34.8 | 0.2 | 300.0 | | |
| PCR06 | STANDARD | | | | ITS1 | 34.7 | 34.8 | 0.2 | 300.0 | | |
| PCR06 | STANDARD | | | | ITS1 | 35.0 | 34.8 | 0.2 | 300.0 | | |
| PCR06 | STANDARD | | | | ITS1 | 39.5 | 39.5 | 0.1 | 30.0 | | |
| PCR06 | STANDARD | | | | ITS1 | 39.6 | 39.5 | 0.1 | 30.0 | | |
| PCR06 | STANDARD | | | | ITS1 | 39.3 | 39.5 | 0.1 | 30.0 | | |

| | | | | | | | | | | | |
|---|---|---|---|---|---|---|---|---|---|---|---|
| PCR06 | sunrise | tank1 | 2 | 0.7 | ITS1 | 28.8 | 28.6 | 0.2 | 11356.9 | 12703.4 | 1170.9 |
| PCR06 | sunrise | tank1 | 2 | 0.7 | ITS1 | 28.5 | 28.6 | 0.2 | 13482.3 | 12703.4 | 1170.9 |
| PCR06 | sunrise | tank1 | 2 | 0.7 | ITS1 | 28.5 | 28.6 | 0.2 | 13271.0 | 12703.4 | 1170.9 |
| PCR06 | sunrise | tank1 | 2 | 2 | ITS1 | 33.7 | 33.7 | 0.3 | 685.1 | 683.0 | 96.6 |
| PCR06 | sunrise | tank1 | 2 | 2 | ITS1 | 34.0 | 33.7 | 0.3 | 585.4 | 683.0 | 96.6 |
| PCR06 | sunrise | tank1 | 2 | 2 | ITS1 | 33.5 | 33.7 | 0.3 | 778.5 | 683.0 | 96.6 |
| PCR06 | sunrise | tank1 | 2 | 8 | ITS1 | 36.9 | 36.4 | 0.5 | 114.9 | 155.0 | 41.4 |
| PCR06 | sunrise | tank1 | 2 | 8 | ITS1 | 36.4 | 36.4 | 0.5 | 152.8 | 155.0 | 41.4 |
| PCR06 | sunrise | tank1 | 2 | 8 | ITS1 | 35.9 | 36.4 | 0.5 | 197.5 | 155.0 | 41.4 |
| PCR06 | sunrise | tank1 | 2 | 20 | ITS1 | 33.8 | 33.7 | 0.2 | 647.3 | 681.2 | 72.0 |
| PCR06 | sunrise | tank1 | 2 | 20 | ITS1 | 33.5 | 33.7 | 0.2 | 763.9 | 681.2 | 72.0 |
| PCR06 | sunrise | tank1 | 2 | 20 | ITS1 | 33.9 | 33.7 | 0.2 | 632.5 | 681.2 | 72.0 |
| PCR06 | sunrise | tank1 | 3 | 0.7 | ITS1 | 28.8 | 28.6 | 0.1 | 11437.2 | 12288.4 | 831.5 |
| PCR06 | sunrise | tank1 | 3 | 0.7 | ITS1 | 28.6 | 28.6 | 0.1 | 12329.6 | 12288.4 | 831.5 |
| PCR06 | sunrise | tank1 | 3 | 0.7 | ITS1 | 28.5 | 28.6 | 0.1 | 13098.5 | 12288.4 | 831.5 |
| PCR06 | sunrise | tank1 | 3 | 2 | ITS1 | 34.0 | 33.9 | 0.1 | 590.9 | 619.0 | 29.0 |
| PCR06 | sunrise | tank1 | 3 | 2 | ITS1 | 33.9 | 33.9 | 0.1 | 617.2 | 619.0 | 29.0 |
| PCR06 | sunrise | tank1 | 3 | 2 | ITS1 | 33.8 | 33.9 | 0.1 | 648.8 | 619.0 | 29.0 |
| PCR06 | sunrise | tank1 | 3 | 8 | ITS1 | 36.7 | 36.5 | 0.3 | 127.9 | 147.0 | 21.7 |
| PCR06 | sunrise | tank1 | 3 | 8 | ITS1 | 36.5 | 36.5 | 0.3 | 142.5 | 147.0 | 21.7 |
| PCR06 | sunrise | tank1 | 3 | 8 | ITS1 | 36.2 | 36.5 | 0.3 | 170.6 | 147.0 | 21.7 |
| PCR06 | sunrise | tank1 | 3 | 20 | ITS1 | 34.0 | 34.0 | 0.1 | 596.0 | 582.6 | 21.0 |
| PCR06 | sunrise | tank1 | 3 | 20 | ITS1 | 34.0 | 34.0 | 0.1 | 593.3 | 582.6 | 21.0 |
| PCR06 | sunrise | tank1 | 3 | 20 | ITS1 | 34.1 | 34.0 | 0.1 | 558.3 | 582.6 | 21.0 |
| PCR06 | sunset | tank1 | 1 | 0.7 | ITS1 | 29.1 | 29.1 | 0.0 | 9326.1 | 9397.0 | 75.9 |
| PCR06 | sunset | tank1 | 1 | 0.7 | ITS1 | 29.1 | 29.1 | 0.0 | 9387.7 | 9397.0 | 75.9 |
| PCR06 | sunset | tank1 | 1 | 0.7 | ITS1 | 29.1 | 29.1 | 0.0 | 9477.1 | 9397.0 | 75.9 |
| PCR06 | sunset | tank1 | 1 | 2 | ITS1 | 35.9 | 35.5 | 0.4 | 203.6 | 261.5 | 59.1 |
| PCR06 | sunset | tank1 | 1 | 2 | ITS1 | 35.1 | 35.5 | 0.4 | 321.8 | 261.5 | 59.1 |
| PCR06 | sunset | tank1 | 1 | 2 | ITS1 | 35.4 | 35.5 | 0.4 | 259.0 | 261.5 | 59.1 |
| PCR06 | sunset | tank1 | 1 | 8 | ITS1 | 36.5 | 36.2 | 0.3 | 143.6 | 169.9 | 31.6 |
| PCR06 | sunset | tank1 | 1 | 8 | ITS1 | 36.3 | 36.2 | 0.3 | 161.0 | 169.9 | 31.6 |
| PCR06 | sunset | tank1 | 1 | 8 | ITS1 | 35.9 | 36.2 | 0.3 | 205.0 | 169.9 | 31.6 |
| PCR06 | sunset | tank1 | 1 | 14 | ITS1 | 37.5 | 37.0 | 0.4 | 82.9 | 107.3 | 22.2 |
| PCR06 | sunset | tank1 | 1 | 14 | ITS1 | 36.7 | 37.0 | 0.4 | 126.4 | 107.3 | 22.2 |
| PCR06 | sunset | tank1 | 1 | 14 | ITS1 | 36.9 | 37.0 | 0.4 | 112.6 | 107.3 | 22.2 |

| | | | | | | | | | | | |
|---|---|---|---|---|---|---|---|---|---|---|---|
| PCR06 | sunrise | tank2 | 3 | 0.7 | ITS1 | 25.3 | 25.2 | 0.1 | 83642.4 | 87763.2 | 4216.9 |
| PCR06 | sunrise | tank2 | 3 | 0.7 | ITS1 | 25.1 | 25.2 | 0.1 | 92070.1 | 87763.2 | 4216.9 |
| PCR06 | sunrise | tank2 | 3 | 0.7 | ITS1 | 25.2 | 25.2 | 0.1 | 87577.2 | 87763.2 | 4216.9 |
| PCR06 | sunrise | tank2 | 3 | 2 | ITS1 | 30.7 | 30.8 | 0.1 | 3728.1 | 3672.1 | 216.0 |
| PCR06 | sunrise | tank2 | 3 | 2 | ITS1 | 30.9 | 30.8 | 0.1 | 3433.6 | 3672.1 | 216.0 |
| PCR06 | sunrise | tank2 | 3 | 2 | ITS1 | 30.7 | 30.8 | 0.1 | 3854.5 | 3672.1 | 216.0 |
| PCR06 | sunrise | tank2 | 3 | 8 | ITS1 | 33.4 | 33.5 | 0.1 | 822.3 | 769.0 | 50.9 |
| PCR06 | sunrise | tank2 | 3 | 8 | ITS1 | 33.5 | 33.5 | 0.1 | 763.8 | 769.0 | 50.9 |
| PCR06 | sunrise | tank2 | 3 | 8 | ITS1 | 33.6 | 33.5 | 0.1 | 720.9 | 769.0 | 50.9 |
| PCR06 | sunrise | tank2 | 3 | 14 | ITS1 | 34.3 | 34.3 | 0.2 | 498.0 | 494.2 | 45.9 |
| PCR06 | sunrise | tank2 | 3 | 14 | ITS1 | 34.5 | 34.3 | 0.2 | 446.5 | 494.2 | 45.9 |
| PCR06 | sunrise | tank2 | 3 | 14 | ITS1 | 34.2 | 34.3 | 0.2 | 538.1 | 494.2 | 45.9 |
| PCR06 | sunrise | tank2 | 3 | 20 | ITS1 | 31.7 | 31.6 | 0.1 | 2176.3 | 2330.8 | 184.3 |
| PCR06 | sunrise | tank2 | 3 | 20 | ITS1 | 31.4 | 31.6 | 0.1 | 2534.8 | 2330.8 | 184.3 |
| PCR06 | sunrise | tank2 | 3 | 20 | ITS1 | 31.6 | 31.6 | 0.1 | 2281.4 | 2330.8 | 184.3 |
| PCR06 | sunset | inlet seawater | | | ITS1 | 39.0 | 39.4 | 0.3 | 33.8 | 27.8 | 5.3 |
| PCR06 | sunset | inlet seawater | | | ITS1 | 39.7 | 39.4 | 0.3 | 23.8 | 27.8 | 5.3 |
| PCR06 | sunset | inlet seawater | | | ITS1 | 39.5 | 39.4 | 0.3 | 25.7 | 27.8 | 5.3 |
| PCR06 | sunrise | inlet seawater | | | ITS1 | 37.4 | 37.7 | 1.2 | 86.9 | 81.1 | 45.5 |
| PCR06 | sunrise | inlet seawater | | | ITS1 | 39.1 | 37.7 | 1.2 | 33.0 | 81.1 | 45.5 |
| PCR06 | sunrise | inlet seawater | | | ITS1 | 36.7 | 37.7 | 1.2 | 123.5 | 81.1 | 45.5 |
| PCR06 | sunset | FilterNC | | | ITS1 | Undetermined | | | | | |
| PCR06 | sunset | FilterNC | | | ITS1 | Undetermined | | | | | |
| PCR06 | sunset | FilterNC | | | ITS1 | Undetermined | | | | | |
| PCR06 | sunrise | FilterNC | | | ITS1 | Undetermined | | | | | |
| PCR06 | sunrise | FilterNC | | | ITS1 | Undetermined | | | | | |
| PCR06 | sunrise | FilterNC | | | ITS1 | Undetermined | | | | | |
| PCR06 | PCRNC | | | | ITS1 | Undetermined | | | | | |
| PCR06 | PCRNC | | | | ITS1 | Undetermined | | | | | |
| PCR06 | PCRNC | | | | ITS1 | Undetermined | | | | | |
| PCR07 | STANDARD | | | | Cyt b | 25.1 | 25.1 | 0.1 | 30000.0 | | |
| PCR07 | STANDARD | | | | Cyt b | 25.0 | 25.1 | 0.1 | 30000.0 | | |
| PCR07 | STANDARD | | | | Cyt b | 25.1 | 25.1 | 0.1 | 30000.0 | | |
| PCR07 | STANDARD | | | | Cyt b | 28.8 | 28.8 | 0.0 | 3000.0 | | |
| PCR07 | STANDARD | | | | Cyt b | 28.7 | 28.8 | 0.0 | 3000.0 | | |
| PCR07 | STANDARD | | | | Cyt b | 28.8 | 28.8 | 0.0 | 3000.0 | | |

| | | | | | | | | | | | |
|---|---|---|---|---|---|---|---|---|---|---|---|
| PCR07 | STANDARD | | | | Cyt b | 32.1 | 32.1 | 0.1 | 300.0 | | |
| PCR07 | STANDARD | | | | Cyt b | 32.3 | 32.1 | 0.1 | 300.0 | | |
| PCR07 | STANDARD | | | | Cyt b | 32.1 | 32.1 | 0.1 | 300.0 | | |
| PCR07 | STANDARD | | | | Cyt b | 35.9 | 35.9 | 0.0 | 30.0 | | |
| PCR07 | STANDARD | | | | Cyt b | 36.0 | 35.9 | 0.0 | 30.0 | | |
| PCR07 | STANDARD | | | | Cyt b | 36.0 | 35.9 | 0.0 | 30.0 | | |
| PCR07 | sunset | tank1 | 2 | 0.7 | Cyt b | 31.1 | 31.0 | 0.1 | 633.1 | 674.0 | 35.6 |
| PCR07 | sunset | tank1 | 2 | 0.7 | Cyt b | 31.0 | 31.0 | 0.1 | 691.1 | 674.0 | 35.6 |
| PCR07 | sunset | tank1 | 2 | 0.7 | Cyt b | 31.0 | 31.0 | 0.1 | 697.8 | 674.0 | 35.6 |
| PCR07 | sunset | tank1 | 2 | 2 | Cyt b | 32.1 | 31.9 | 0.1 | 346.4 | 375.3 | 29.6 |
| PCR07 | sunset | tank1 | 2 | 2 | Cyt b | 31.9 | 31.9 | 0.1 | 373.9 | 375.3 | 29.6 |
| PCR07 | sunset | tank1 | 2 | 2 | Cyt b | 31.8 | 31.9 | 0.1 | 405.6 | 375.3 | 29.6 |
| PCR07 | sunset | tank1 | 2 | 8 | Cyt b | 34.4 | 34.5 | 0.1 | 76.2 | 71.8 | 4.5 |
| PCR07 | sunset | tank1 | 2 | 8 | Cyt b | 34.5 | 34.5 | 0.1 | 72.0 | 71.8 | 4.5 |
| PCR07 | sunset | tank1 | 2 | 8 | Cyt b | 34.6 | 34.5 | 0.1 | 67.3 | 71.8 | 4.5 |
| PCR07 | sunset | tank1 | 2 | 14 | Cyt b | 36.0 | 36.1 | 0.1 | 27.5 | 27.0 | 2.1 |
| PCR07 | sunset | tank1 | 2 | 14 | Cyt b | 35.9 | 36.1 | 0.1 | 28.9 | 27.0 | 2.1 |
| PCR07 | sunset | tank1 | 2 | 14 | Cyt b | 36.2 | 36.1 | 0.1 | 24.7 | 27.0 | 2.1 |
| PCR07 | sunset | tank1 | 2 | 20 | Cyt b | 33.7 | 33.7 | 0.2 | 118.7 | 124.3 | 16.2 |
| PCR07 | sunset | tank1 | 2 | 20 | Cyt b | 33.8 | 33.7 | 0.2 | 111.7 | 124.3 | 16.2 |
| PCR07 | sunset | tank1 | 2 | 20 | Cyt b | 33.5 | 33.7 | 0.2 | 142.5 | 124.3 | 16.2 |
| PCR07 | sunset | tank1 | 3 | 0.7 | Cyt b | 31.0 | 30.9 | 0.1 | 688.5 | 748.8 | 71.6 |
| PCR07 | sunset | tank1 | 3 | 0.7 | Cyt b | 30.9 | 30.9 | 0.1 | 730.0 | 748.8 | 71.6 |
| PCR07 | sunset | tank1 | 3 | 0.7 | Cyt b | 30.7 | 30.9 | 0.1 | 828.0 | 748.8 | 71.6 |
| PCR07 | sunset | tank1 | 3 | 2 | Cyt b | 31.7 | 31.7 | 0.1 | 438.1 | 424.0 | 17.0 |
| PCR07 | sunset | tank1 | 3 | 2 | Cyt b | 31.7 | 31.7 | 0.1 | 428.7 | 424.0 | 17.0 |
| PCR07 | sunset | tank1 | 3 | 2 | Cyt b | 31.8 | 31.7 | 0.1 | 405.2 | 424.0 | 17.0 |
| PCR07 | sunset | tank1 | 3 | 8 | Cyt b | 35.6 | 35.3 | 0.3 | 35.8 | 43.9 | 7.9 |
| PCR07 | sunset | tank1 | 3 | 8 | Cyt b | 35.3 | 35.3 | 0.3 | 44.4 | 43.9 | 7.9 |
| PCR07 | sunset | tank1 | 3 | 8 | Cyt b | 35.0 | 35.3 | 0.3 | 51.5 | 43.9 | 7.9 |
| PCR07 | sunset | tank1 | 3 | 14 | Cyt b | 35.0 | 35.4 | 0.5 | 54.5 | 43.0 | 13.4 |
| PCR07 | sunset | tank1 | 3 | 14 | Cyt b | 35.2 | 35.4 | 0.5 | 46.2 | 43.0 | 13.4 |
| PCR07 | sunset | tank1 | 3 | 14 | Cyt b | 36.0 | 35.4 | 0.5 | 28.3 | 43.0 | 13.4 |
| PCR07 | sunset | tank1 | 3 | 20 | Cyt b | 34.5 | 34.4 | 0.5 | 74.0 | 78.3 | 23.8 |
| PCR07 | sunset | tank1 | 3 | 20 | Cyt b | 34.9 | 34.4 | 0.5 | 56.9 | 78.3 | 23.8 |
| PCR07 | sunset | tank1 | 3 | 20 | Cyt b | 33.9 | 34.4 | 0.5 | 104.0 | 78.3 | 23.8 |

| | | | | | | | | | | | |
|---|---|---|---|---|---|---|---|---|---|---|---|
| PCR07 | sunset | tank2 | 1 | 0.7 | Cyt b | 28.5 | 28.4 | 0.1 | 3418.0 | 3668.5 | 226.8 |
| PCR07 | sunset | tank2 | 1 | 0.7 | Cyt b | 28.3 | 28.4 | 0.1 | 3727.2 | 3668.5 | 226.8 |
| PCR07 | sunset | tank2 | 1 | 0.7 | Cyt b | 28.3 | 28.4 | 0.1 | 3860.1 | 3668.5 | 226.8 |
| PCR07 | sunset | tank2 | 1 | 2 | Cyt b | 31.4 | 31.5 | 0.1 | 522.0 | 504.7 | 17.4 |
| PCR07 | sunset | tank2 | 1 | 2 | Cyt b | 31.5 | 31.5 | 0.1 | 505.0 | 504.7 | 17.4 |
| PCR07 | sunset | tank2 | 1 | 2 | Cyt b | 31.5 | 31.5 | 0.1 | 487.1 | 504.7 | 17.4 |
| PCR07 | sunset | tank2 | 1 | 8 | Cyt b | 33.0 | 32.9 | 0.1 | 193.0 | 202.6 | 10.7 |
| PCR07 | sunset | tank2 | 1 | 8 | Cyt b | 32.8 | 32.9 | 0.1 | 214.1 | 202.6 | 10.7 |
| PCR07 | sunset | tank2 | 1 | 8 | Cyt b | 32.9 | 32.9 | 0.1 | 200.6 | 202.6 | 10.7 |
| PCR07 | sunset | tank2 | 1 | 14 | Cyt b | 33.0 | 32.9 | 0.2 | 191.1 | 205.9 | 25.3 |
| PCR07 | sunset | tank2 | 1 | 14 | Cyt b | 32.7 | 32.9 | 0.2 | 235.1 | 205.9 | 25.3 |
| PCR07 | sunset | tank2 | 1 | 14 | Cyt b | 33.0 | 32.9 | 0.2 | 191.5 | 205.9 | 25.3 |
| PCR07 | sunset | tank2 | 1 | 20 | Cyt b | 30.8 | 30.7 | 0.1 | 775.7 | 818.5 | 52.4 |
| PCR07 | sunset | tank2 | 1 | 20 | Cyt b | 30.6 | 30.7 | 0.1 | 877.0 | 818.5 | 52.4 |
| PCR07 | sunset | tank2 | 1 | 20 | Cyt b | 30.7 | 30.7 | 0.1 | 802.9 | 818.5 | 52.4 |
| PCR07 | sunset | tank2 | 2 | 0.7 | Cyt b | 29.0 | 29.1 | 0.1 | 2375.5 | 2354.7 | 76.7 |
| PCR07 | sunset | tank2 | 2 | 0.7 | Cyt b | 29.0 | 29.1 | 0.1 | 2418.8 | 2354.7 | 76.7 |
| PCR07 | sunset | tank2 | 2 | 0.7 | Cyt b | 29.1 | 29.1 | 0.1 | 2269.7 | 2354.7 | 76.7 |
| PCR07 | sunset | tank2 | 2 | 2 | Cyt b | 30.7 | 30.6 | 0.2 | 827.5 | 896.1 | 93.3 |
| PCR07 | sunset | tank2 | 2 | 2 | Cyt b | 30.6 | 30.6 | 0.2 | 858.4 | 896.1 | 93.3 |
| PCR07 | sunset | tank2 | 2 | 2 | Cyt b | 30.4 | 30.6 | 0.2 | 1002.3 | 896.1 | 93.3 |
| PCR07 | sunset | tank2 | 2 | 8 | Cyt b | 32.6 | 32.4 | 0.3 | 240.3 | 286.4 | 49.1 |
| PCR07 | sunset | tank2 | 2 | 8 | Cyt b | 32.4 | 32.4 | 0.3 | 280.7 | 286.4 | 49.1 |
| PCR07 | sunset | tank2 | 2 | 8 | Cyt b | 32.1 | 32.4 | 0.3 | 338.1 | 286.4 | 49.1 |
| PCR07 | sunset | tank2 | 2 | 14 | Cyt b | 33.4 | 33.2 | 0.2 | 151.4 | 169.1 | 20.8 |
| PCR07 | sunset | tank2 | 2 | 14 | Cyt b | 33.0 | 33.2 | 0.2 | 192.1 | 169.1 | 20.8 |
| PCR07 | sunset | tank2 | 2 | 14 | Cyt b | 33.2 | 33.2 | 0.2 | 163.8 | 169.1 | 20.8 |
| PCR07 | sunset | tank2 | 2 | 20 | Cyt b | 34.7 | 34.6 | 0.2 | 62.6 | 69.1 | 8.7 |
| PCR07 | sunset | tank2 | 2 | 20 | Cyt b | 34.4 | 34.6 | 0.2 | 79.1 | 69.1 | 8.7 |
| PCR07 | sunset | tank2 | 2 | 20 | Cyt b | 34.7 | 34.6 | 0.2 | 65.7 | 69.1 | 8.7 |
| PCR07 | sunset | tank2 | 3 | 0.7 | Cyt b | 28.7 | 28.6 | 0.1 | 3032.6 | 3266.8 | 211.2 |
| PCR07 | sunset | tank2 | 3 | 0.7 | Cyt b | 28.5 | 28.6 | 0.1 | 3324.9 | 3266.8 | 211.2 |
| PCR07 | sunset | tank2 | 3 | 0.7 | Cyt b | 28.5 | 28.6 | 0.1 | 3442.9 | 3266.8 | 211.2 |
| PCR07 | sunset | tank2 | 3 | 2 | Cyt b | 30.1 | 30.0 | 0.1 | 1212.8 | 1309.9 | 84.1 |
| PCR07 | sunset | tank2 | 3 | 2 | Cyt b | 29.9 | 30.0 | 0.1 | 1361.4 | 1309.9 | 84.1 |
| PCR07 | sunset | tank2 | 3 | 2 | Cyt b | 29.9 | 30.0 | 0.1 | 1355.6 | 1309.9 | 84.1 |

| | | | | | | | | | | | |
|---|---|---|---|---|---|---|---|---|---|---|---|
| PCR07 | sunset | tank2 | 3 | 8 | Cyt b | 31.2 | 31.1 | 0.1 | 598.9 | 644.0 | 46.7 |
| PCR07 | sunset | tank2 | 3 | 8 | Cyt b | 31.0 | 31.1 | 0.1 | 692.1 | 644.0 | 46.7 |
| PCR07 | sunset | tank2 | 3 | 8 | Cyt b | 31.1 | 31.1 | 0.1 | 640.8 | 644.0 | 46.7 |
| PCR07 | sunset | tank2 | 3 | 14 | Cyt b | 34.8 | 34.8 | 0.0 | 61.1 | 61.5 | 1.4 |
| PCR07 | sunset | tank2 | 3 | 14 | Cyt b | 34.7 | 34.8 | 0.0 | 63.1 | 61.5 | 1.4 |
| PCR07 | sunset | tank2 | 3 | 14 | Cyt b | 34.8 | 34.8 | 0.0 | 60.4 | 61.5 | 1.4 |
| PCR07 | sunset | tank2 | 3 | 20 | Cyt b | 33.2 | 33.3 | 0.2 | 170.6 | 162.3 | 18.8 |
| PCR07 | sunset | tank2 | 3 | 20 | Cyt b | 33.5 | 33.3 | 0.2 | 140.9 | 162.3 | 18.8 |
| PCR07 | sunset | tank2 | 3 | 20 | Cyt b | 33.1 | 33.3 | 0.2 | 175.6 | 162.3 | 18.8 |
| PCR07 | PCRNC | | | | Cyt b | Undetermined | | | | | |
| PCR07 | PCRNC | | | | Cyt b | Undetermined | | | | | |
| PCR07 | PCRNC | | | | Cyt b | Undetermined | | | | | |
| PCR08 | STANDARD | | | | ITS1 | 26.5 | 26.3 | 0.2 | 30000.0 | | |
| PCR08 | STANDARD | | | | ITS1 | 26.2 | 26.3 | 0.2 | 30000.0 | | |
| PCR08 | STANDARD | | | | ITS1 | 26.3 | 26.3 | 0.2 | 30000.0 | | |
| PCR08 | STANDARD | | | | ITS1 | 30.5 | 30.4 | 0.1 | 3000.0 | | |
| PCR08 | STANDARD | | | | ITS1 | 30.3 | 30.4 | 0.1 | 3000.0 | | |
| PCR08 | STANDARD | | | | ITS1 | 30.4 | 30.4 | 0.1 | 3000.0 | | |
| PCR08 | STANDARD | | | | ITS1 | 33.8 | 33.9 | 0.1 | 300.0 | | |
| PCR08 | STANDARD | | | | ITS1 | 34.0 | 33.9 | 0.1 | 300.0 | | |
| PCR08 | STANDARD | | | | ITS1 | 33.8 | 33.9 | 0.1 | 300.0 | | |
| PCR08 | STANDARD | | | | ITS1 | 38.6 | 38.3 | 0.4 | 30.0 | | |
| PCR08 | STANDARD | | | | ITS1 | 37.9 | 38.3 | 0.4 | 30.0 | | |
| PCR08 | STANDARD | | | | ITS1 | 38.3 | 38.3 | 0.4 | 30.0 | | |
| PCR08 | sunset | tank1 | 2 | 0.7 | ITS1 | 28.6 | 28.5 | 0.1 | 7841.2 | 8223.5 | 473.0 |
| PCR08 | sunset | tank1 | 2 | 0.7 | ITS1 | 28.6 | 28.5 | 0.1 | 8076.8 | 8223.5 | 473.0 |
| PCR08 | sunset | tank1 | 2 | 0.7 | ITS1 | 28.4 | 28.5 | 0.1 | 8752.5 | 8223.5 | 473.0 |
| PCR08 | sunset | tank1 | 2 | 2 | ITS1 | 35.4 | 35.2 | 0.3 | 152.4 | 169.4 | 26.2 |
| PCR08 | sunset | tank1 | 2 | 2 | ITS1 | 35.3 | 35.2 | 0.3 | 156.3 | 169.4 | 26.2 |
| PCR08 | sunset | tank1 | 2 | 2 | ITS1 | 34.9 | 35.2 | 0.3 | 199.6 | 169.4 | 26.2 |
| PCR08 | sunset | tank1 | 2 | 8 | ITS1 | 35.0 | 35.0 | 0.1 | 190.1 | 191.2 | 10.4 |
| PCR08 | sunset | tank1 | 2 | 8 | ITS1 | 34.9 | 35.0 | 0.1 | 202.1 | 191.2 | 10.4 |
| PCR08 | sunset | tank1 | 2 | 8 | ITS1 | 35.1 | 35.0 | 0.1 | 181.4 | 191.2 | 10.4 |
| PCR08 | sunset | tank1 | 2 | 20 | ITS1 | 36.5 | 37.0 | 0.6 | 77.6 | 58.6 | 18.9 |
| PCR08 | sunset | tank1 | 2 | 20 | ITS1 | 37.6 | 37.0 | 0.6 | 39.8 | 58.6 | 18.9 |
| PCR08 | sunset | tank1 | 2 | 20 | ITS1 | 37.0 | 37.0 | 0.6 | 58.5 | 58.6 | 18.9 |

| | | | | | | | | | | | |
|---|---|---|---|---|---|---|---|---|---|---|---|
| PCR08 | sunset | tank1 | 3 | 0.7 | ITS1 | 28.6 | 28.5 | 0.1 | 8045.7 | 8605.2 | 490.8 |
| PCR08 | sunset | tank1 | 3 | 0.7 | ITS1 | 28.4 | 28.5 | 0.1 | 8806.7 | 8605.2 | 490.8 |
| PCR08 | sunset | tank1 | 3 | 0.7 | ITS1 | 28.4 | 28.5 | 0.1 | 8963.2 | 8605.2 | 490.8 |
| PCR08 | sunset | tank1 | 3 | 8 | ITS1 | 37.2 | 36.7 | 0.5 | 50.6 | 73.2 | 19.6 |
| PCR08 | sunset | tank1 | 3 | 8 | ITS1 | 36.4 | 36.7 | 0.5 | 83.0 | 73.2 | 19.6 |
| PCR08 | sunset | tank1 | 3 | 8 | ITS1 | 36.3 | 36.7 | 0.5 | 86.0 | 73.2 | 19.6 |
| PCR08 | sunset | tank1 | 3 | 14 | ITS1 | 37.0 | 37.0 | 0.3 | 56.9 | 58.9 | 9.8 |
| PCR08 | sunset | tank1 | 3 | 14 | ITS1 | 37.3 | 37.0 | 0.3 | 50.1 | 58.9 | 9.8 |
| PCR08 | sunset | tank1 | 3 | 14 | ITS1 | 36.7 | 37.0 | 0.3 | 69.5 | 58.9 | 9.8 |
| PCR08 | sunset | tank2 | 1 | 0.7 | ITS1 | 25.1 | 25.0 | 0.1 | 62102.8 | 64369.7 | 2061.5 |
| PCR08 | sunset | tank2 | 1 | 0.7 | ITS1 | 25.0 | 25.0 | 0.1 | 66132.1 | 64369.7 | 2061.5 |
| PCR08 | sunset | tank2 | 1 | 0.7 | ITS1 | 25.0 | 25.0 | 0.1 | 64874.2 | 64369.7 | 2061.5 |
| PCR08 | sunset | tank2 | 1 | 2 | ITS1 | 32.9 | 33.2 | 0.3 | 625.8 | 540.9 | 98.7 |
| PCR08 | sunset | tank2 | 1 | 2 | ITS1 | 33.6 | 33.2 | 0.3 | 432.6 | 540.9 | 98.7 |
| PCR08 | sunset | tank2 | 1 | 2 | ITS1 | 33.1 | 33.2 | 0.3 | 564.1 | 540.9 | 98.7 |
| PCR08 | sunset | tank2 | 1 | 8 | ITS1 | 32.3 | 32.2 | 0.1 | 918.8 | 989.4 | 73.2 |
| PCR08 | sunset | tank2 | 1 | 8 | ITS1 | 32.2 | 32.2 | 0.1 | 984.6 | 989.4 | 73.2 |
| PCR08 | sunset | tank2 | 1 | 8 | ITS1 | 32.0 | 32.2 | 0.1 | 1065.0 | 989.4 | 73.2 |
| PCR08 | sunset | tank2 | 1 | 14 | ITS1 | 32.9 | 33.0 | 0.1 | 653.1 | 615.9 | 32.9 |
| PCR08 | sunset | tank2 | 1 | 14 | ITS1 | 33.0 | 33.0 | 0.1 | 604.2 | 615.9 | 32.9 |
| PCR08 | sunset | tank2 | 1 | 14 | ITS1 | 33.0 | 33.0 | 0.1 | 590.5 | 615.9 | 32.9 |
| PCR08 | sunset | tank2 | 1 | 20 | ITS1 | 31.7 | 31.6 | 0.1 | 1306.0 | 1364.7 | 52.8 |
| PCR08 | sunset | tank2 | 1 | 20 | ITS1 | 31.6 | 31.6 | 0.1 | 1379.5 | 1364.7 | 52.8 |
| PCR08 | sunset | tank2 | 1 | 20 | ITS1 | 31.6 | 31.6 | 0.1 | 1408.4 | 1364.7 | 52.8 |
| PCR08 | sunset | tank2 | 2 | 0.7 | ITS1 | 25.9 | 25.8 | 0.0 | 38870.8 | 40211.3 | 1166.6 |
| PCR08 | sunset | tank2 | 2 | 0.7 | ITS1 | 25.8 | 25.8 | 0.0 | 40997.5 | 40211.3 | 1166.6 |
| PCR08 | sunset | tank2 | 2 | 0.7 | ITS1 | 25.8 | 25.8 | 0.0 | 40765.5 | 40211.3 | 1166.6 |
| PCR08 | sunset | tank2 | 2 | 2 | ITS1 | 31.6 | 31.8 | 0.2 | 1338.7 | 1200.9 | 121.8 |
| PCR08 | sunset | tank2 | 2 | 2 | ITS1 | 32.0 | 31.8 | 0.2 | 1107.8 | 1200.9 | 121.8 |
| PCR08 | sunset | tank2 | 2 | 2 | ITS1 | 31.9 | 31.8 | 0.2 | 1156.2 | 1200.9 | 121.8 |
| PCR08 | sunset | tank2 | 2 | 8 | ITS1 | 32.7 | 32.6 | 0.1 | 729.8 | 745.8 | 37.2 |
| PCR08 | sunset | tank2 | 2 | 8 | ITS1 | 32.7 | 32.6 | 0.1 | 719.2 | 745.8 | 37.2 |
| PCR08 | sunset | tank2 | 2 | 8 | ITS1 | 32.5 | 32.6 | 0.1 | 788.3 | 745.8 | 37.2 |
| PCR08 | sunset | tank2 | 2 | 14 | ITS1 | 34.0 | 33.8 | 0.3 | 334.8 | 378.0 | 57.6 |
| PCR08 | sunset | tank2 | 2 | 14 | ITS1 | 33.5 | 33.8 | 0.3 | 443.3 | 378.0 | 57.6 |
| PCR08 | sunset | tank2 | 2 | 14 | ITS1 | 33.9 | 33.8 | 0.3 | 355.8 | 378.0 | 57.6 |

| | | | | | | | | | | | |
|---|---|---|---|---|---|---|---|---|---|---|---|
| PCR08 | sunset | tank2 | 2 | 20 | ITS1 | 36.5 | 36.0 | 0.4 | 78.5 | 107.8 | 25.4 |
| PCR08 | sunset | tank2 | 2 | 20 | ITS1 | 35.7 | 36.0 | 0.4 | 122.8 | 107.8 | 25.4 |
| PCR08 | sunset | tank2 | 2 | 20 | ITS1 | 35.7 | 36.0 | 0.4 | 122.1 | 107.8 | 25.4 |
| PCR08 | sunset | tank2 | 3 | 0.7 | ITS1 | 25.7 | 25.4 | 0.2 | 44864.0 | 51806.5 | 6333.4 |
| PCR08 | sunset | tank2 | 3 | 0.7 | ITS1 | 25.4 | 25.4 | 0.2 | 53287.0 | 51806.5 | 6333.4 |
| PCR08 | sunset | tank2 | 3 | 0.7 | ITS1 | 25.2 | 25.4 | 0.2 | 57268.5 | 51806.5 | 6333.4 |
| PCR08 | sunset | tank2 | 3 | 2 | ITS1 | 33.7 | 33.4 | 0.3 | 400.5 | 497.3 | 83.8 |
| PCR08 | sunset | tank2 | 3 | 2 | ITS1 | 33.2 | 33.4 | 0.3 | 544.7 | 497.3 | 83.8 |
| PCR08 | sunset | tank2 | 3 | 2 | ITS1 | 33.2 | 33.4 | 0.3 | 546.7 | 497.3 | 83.8 |
| PCR08 | sunset | tank2 | 3 | 8 | ITS1 | 32.6 | 32.5 | 0.1 | 755.0 | 798.0 | 63.7 |
| PCR08 | sunset | tank2 | 3 | 8 | ITS1 | 32.6 | 32.5 | 0.1 | 767.8 | 798.0 | 63.7 |
| PCR08 | sunset | tank2 | 3 | 8 | ITS1 | 32.4 | 32.5 | 0.1 | 871.1 | 798.0 | 63.7 |
| PCR08 | sunset | tank2 | 3 | 14 | ITS1 | 35.3 | 35.5 | 0.4 | 159.8 | 142.0 | 30.5 |
| PCR08 | sunset | tank2 | 3 | 14 | ITS1 | 36.0 | 35.5 | 0.4 | 106.7 | 142.0 | 30.5 |
| PCR08 | sunset | tank2 | 3 | 14 | ITS1 | 35.3 | 35.5 | 0.4 | 159.3 | 142.0 | 30.5 |
| PCR08 | sunset | tank2 | 3 | 20 | ITS1 | 34.7 | 34.8 | 0.1 | 221.7 | 206.0 | 14.9 |
| PCR08 | sunset | tank2 | 3 | 20 | ITS1 | 35.0 | 34.8 | 0.1 | 192.0 | 206.0 | 14.9 |
| PCR08 | sunset | tank2 | 3 | 20 | ITS1 | 34.9 | 34.8 | 0.1 | 204.3 | 206.0 | 14.9 |
| PCR08 | PCRNC | | | | ITS1 | Undetermined | | | | | |
| PCR08 | PCRNC | | | | ITS1 | Undetermined | | | | | |
| PCR08 | PCRNC | | | | ITS1 | Undetermined | | | | | |
| PCR09 | STANDARD | | | | ITS | 25.1 | 24.8 | 0.2 | 30000.0 | | |
| PCR09 | STANDARD | | | | ITS | 24.7 | 24.8 | 0.2 | 30000.0 | | |
| PCR09 | STANDARD | | | | ITS | 24.7 | 24.8 | 0.2 | 30000.0 | | |
| PCR09 | STANDARD | | | | ITS | 28.9 | 28.8 | 0.1 | 3000.0 | | |
| PCR09 | STANDARD | | | | ITS | 28.8 | 28.8 | 0.1 | 3000.0 | | |
| PCR09 | STANDARD | | | | ITS | 28.8 | 28.8 | 0.1 | 3000.0 | | |
| PCR09 | STANDARD | | | | ITS | 32.4 | 32.6 | 0.1 | 300.0 | | |
| PCR09 | STANDARD | | | | ITS | 32.7 | 32.6 | 0.1 | 300.0 | | |
| PCR09 | STANDARD | | | | ITS | 32.6 | 32.6 | 0.1 | 300.0 | | |
| PCR09 | STANDARD | | | | ITS | 36.2 | 36.2 | 0.6 | 30.0 | | |
| PCR09 | STANDARD | | | | ITS | 35.5 | 36.2 | 0.6 | 30.0 | | |
| PCR09 | STANDARD | | | | ITS | NA | | | | | |
| PCR09 | sunrise | tank1 | 2 | 14 | ITS | 36.7 | 36.9 | 0.7 | 22.2 | 21.6 | 8.1 |
| PCR09 | sunrise | tank1 | 2 | 14 | ITS | 37.6 | 36.9 | 0.7 | 13.2 | 21.6 | 8.1 |
| PCR09 | sunrise | tank1 | 2 | 14 | ITS | 36.3 | 36.9 | 0.7 | 29.3 | 21.6 | 8.1 |

| | | | | | | | | | | | |
|---|---|---|---|---|---|---|---|---|---|---|---|
| PCR09 | sunrise | tank1 | 3 | 14 | ITS | 37.8 | 38.4 | 1.6 | 11.3 | 10.1 | 7.0 |
| PCR09 | sunrise | tank1 | 3 | 14 | ITS | 40.2 | 38.4 | 1.6 | 2.6 | 10.1 | 7.0 |
| PCR09 | sunrise | tank1 | 3 | 14 | ITS | 37.2 | 38.4 | 1.6 | 16.4 | 10.1 | 7.0 |
| PCR09 | sunset | tank1 | 1 | 20 | ITS | 35.3 | 35.3 | 0.2 | 52.5 | 53.4 | 6.6 |
| PCR09 | sunset | tank1 | 1 | 20 | ITS | 35.5 | 35.3 | 0.2 | 47.3 | 53.4 | 6.6 |
| PCR09 | sunset | tank1 | 1 | 20 | ITS | 35.1 | 35.3 | 0.2 | 60.3 | 53.4 | 6.6 |
| PCR09 | sunset | tank1 | 2 | 14 | ITS | 35.8 | 35.5 | 0.2 | 40.1 | 46.8 | 5.8 |
| PCR09 | sunset | tank1 | 2 | 14 | ITS | 35.4 | 35.5 | 0.2 | 49.8 | 46.8 | 5.8 |
| PCR09 | sunset | tank1 | 2 | 14 | ITS | 35.4 | 35.5 | 0.2 | 50.4 | 46.8 | 5.8 |
| PCR09 | sunset | tank1 | 3 | 2 | ITS | 33.3 | 33.2 | 0.0 | 186.3 | 188.4 | 3.3 |
| PCR09 | sunset | tank1 | 3 | 2 | ITS | 33.2 | 33.2 | 0.0 | 192.2 | 188.4 | 3.3 |
| PCR09 | sunset | tank1 | 3 | 2 | ITS | 33.3 | 33.2 | 0.0 | 186.8 | 188.4 | 3.3 |
| PCR09 | sunset | tank1 | 3 | 20 | ITS | 35.2 | 35.3 | 0.3 | 58.2 | 54.4 | 8.4 |
| PCR09 | sunset | tank1 | 3 | 20 | ITS | 35.6 | 35.3 | 0.3 | 44.8 | 54.4 | 8.4 |
| PCR09 | sunset | tank1 | 3 | 20 | ITS | 35.1 | 35.3 | 0.3 | 60.1 | 54.4 | 8.4 |
| PCR09 | PCRNC | | | | ITS | Undetermined | | | | | |
| PCR09 | PCRNC | | | | ITS | Undetermined | | | | | |
| PCR09 | PCRNC | | | | ITS | Undetermined | | | | | |
